# Gene regulatory divergence underlies tissue-specific and sex-specific misexpression in interspecies nematode hybrids

**DOI:** 10.64898/2026.06.24.734386

**Authors:** Athmaja Viswanath, Daniel D. Fusca, John A. Calarco, Asher D. Cutter

## Abstract

Gene regulatory divergence has emerged as a key feature in speciation, influencing gene expression differences that accumulate between diverging populations. Transcriptional regulation, mediated by *cis*- and *trans*-acting factors, modulates diverse developmental processes and is responsible for distinct species-specific gene expression profiles. Within interspecies hybrid individuals, negative interactions between divergent *cis*- and *trans*-acting factors can lead to gene misregulation and hybrid dysfunction at the organismal level. Such gene regulatory mismatch might disproportionately impact sex-biased and tissue-biased gene regulatory networks due to their unique selective pressures. To address these issues, we investigated the role of regulatory divergence in asymmetric hybrid incompatibility between sister species of *Caenorhabditis* nematodes (*C. remanei, C. latens*) by analyzing gene expression of reciprocal hybrids for each sex and key tissue types. Despite severe hybrid male sterility, hybrid males showed less misexpression of sex-biased genes than hybrid females, suggesting that the organismal phenotypic outputs of male-biased gene regulatory networks are more vulnerable to disruption than female-biased genetic networks. Additionally, we found more genes associated with *cis*- than *trans*-regulatory divergence, supporting the notion of a disproportionate role for *cis*-regulatory divergence between species. Moreover, we document extensive *cis*-*trans* compensatory X-linked regulatory divergence specifically from male transcriptomes, indicating distinct molecular evolutionary outcomes of stabilizing selection on regulatory controls in males and females. These insights derived from asymmetric hybrid misexpression expand our understanding of the evolution sex-biased gene regulation in the face of stabilizing selection and identify candidate genes contributing to *Caenorhabditis* post-zygotic reproductive isolation.

## Introduction

Reproductive isolation is an essential feature of speciation that prevents gene flow between diverging populations and maintains species boundaries. Unfit offspring between species are a direct consequence of genetically intrinsic postzygotic barriers, which arise due to negative interactions between genetic factors from different species within the hybrid genome. Such negative epistatic genetic interactions are commonly referred to as Dobzhansky-Muller incompatibilities (DMIs) (Bateson 1909; Dobzhansky 1937; Muller 1942), which can hinder the normal progression of the hybrid’s developmental processes. Disrupted post-zygotic development ultimately results in hybrid dysfunction that can cause extreme phenotypes such as inviability and sterility (Coyne & Allen Orr, 1998; Orr, 1996), as well as more subtle defects such as delayed development (Bomblies et al., 2007; Matute & Coyne, 2010) and behavioral abnormalities (Rice & McQuillan, 2018; Turissini et al., 2017). A prominent pattern observed in F1 interspecies hybrids, known as Haldane’s Rule, is the greater susceptibility of the heterogametic sex to hybrid dysfunction (Haldane, 1922) and is supported by extensive evidence in diverse taxonomic groups (Cowell, 2022; Coyne & Orr, 2004; Delph & Demuth, 2016; Schluter & Rieseberg, 2022; Cutter, 2024). Numerous hypotheses are proposed to account for the underlying mechanism for such sexually asymmetric hybrid dysfunction, including the dominance theory, which posits that sex-asymmetries are attributable to X- or Z-linked genes in the heterogametic sex, the faster male theory, which suggests that dysfunction may arise from the rapid divergence of male-biased genes, and the faster-X theory, which proposes that rapid divergence of X-linked loci may result in X-linked incompatibilities, among others (Turelli, 1998; Presgraves, 2010; Delph & Demuth, 2016; Cowell, 2022; Cutter, 2024). Another asymmetry pertaining to interspecies hybrids is Darwin’s corollary to Haldane’s rule (Turelli & Moyle, 2007), which highlights that the severity of hybrid dysfunction often is sensitive to the specific direction of the cross (i.e. maternal parent species) due to uniparentally inherited factors that contribute to unidirectional incompatibilities, such as X-linked and mitochondrial genes, and epigenetic factors (Turelli & Moyle, 2007). Despite the considerable efforts made in studying the genetic basis of post-zygotic reproductive isolation for almost a century, and the aforementioned broad trends, a consensus regarding the relative importance of specific causal mechanisms across different taxa has not yet been reached.

Divergence in the mechanisms that regulate gene expression has been shown to play an important role in maintaining differences between species and variations within populations (Mack & Nachman, 2017; Runemark et al., 2024; Signor & Nuzhdin, 2018; Wittkopp et al., 2004). Gene regulatory mechanisms can govern the quantity, location, functioning, and interactions of genes, thus dictating crucial developmental processes in organisms and upholding species-specific distinctions in expression (Mack & Nachman, 2017; Signor & Nuzhdin, 2018). Gene regulation at the transcriptional level relies on the interplay between *cis*-acting elements (e.g., promoter or enhancer sequences) and *trans*-acting factors (e.g., transcription factors), with some *trans*-acting factors capable of operating post-transcriptionally (e.g., miRNAs). *cis*-regulatory elements are commonly located upstream of genes and primarily impact gene expression in a localized manner, whereas *trans*-acting factors arise from other regions of the genome and typically can exert control over the regulation of multiple target genes (Mack & Nachman, 2017). In F1 interspecies hybrids, interactions between divergent *cis-* and *trans*-acting factors from different parental species can lead to gene misregulation, impacting developmental pathways and resulting in hybrid incompatibilities like inviability and sterility (Mack & Nachman, 2017). The relative influence of *trans*-regulatory mutations on gene expression is expected to be a greater source of variation within a species than between species, potentially due to their increased pleiotropic effects as they impact multiple targets, to then get weeded out or compensated over time (Metzger et al., 2017; Signor & Nuzhdin, 2018). Changes in *cis*-regulatory elements are expected to play a comparable role to *trans*-acting factors (Coolon et al., 2015; Gomes & Civetta, 2015; Guerrero et al., 2016; McManus et al., 2010; Shi et al., 2012), or a stronger role (Mack et al., 2016; Wittkopp & Kalay, 2012), in creating gene expression differences between species (Carroll, 2005; Hill et al., 2021; Stern & Orgogozo, 2009).

Beyond simple divergence of regulatory features, *cis-* and *trans*-acting factors can co-evolve to compensate for previous evolutionary changes and maintain profiles of gene expression at similar phenotypic levels between species. Such compensatory changes that underlie the expression phenotype are expected to evolve under stabilizing selection and often result in developmental system drift, a phenomenon whereby gene expression phenotype is maintained between species despite molecular evolution of gene regulatory sequences (Cutter & Bundus, 2020; Mack & Nachman, 2017; Schiffman & Ralph, 2022; Signor & Nuzhdin, 2018; True & Haag, 2001). Coevolved regulatory changes, however, can break down in F1 hybrids and manifest as hybrid misexpression due to dysfunctional *cis*-by-*trans* interactions (Mack & Nachman, 2017). Divergence in such gene regulatory mechanisms appears to play a key role in reproductive isolation, with evidence across different taxonomic groups, although debate continues about the relative contributions of alternative molecular mechanisms (Guerrero et al., 2016; X. C. Li & Fay, 2017; Mack et al., 2016; Mack & Nachman, 2017; McManus et al., 2010; Meiklejohn et al., 2014; Sánchez-Ramírez et al., 2021).

Gene regulatory mechanisms also modulate gene expression of the same genes in tissue-biased and sex-biased manners for distinct cell types or male versus female individuals (Ellegren & Parsch, 2007). Genes with such context-dependent expression can experience distinct selective pressures, and might contribute disproportionately to species differences due to their potentially faster evolution than genes with non-differential expression (Assis et al., 2012; Harrison et al., 2015; Parisi et al., 2004; Viswanath et al., 2025). Misregulation of sex-biased genes, in particular, can manifest as dysfunction in sex-specific developmental programs in hybrids to influence sex-specific hybrid incompatibility and Haldane’s rule (Assis et al., 2012; Cutter, 2023a; Gomes & Civetta, 2015; Ranz et al., 2004; Sánchez-Ramírez et al., 2021). Despite such plausible connections between hybrid sterility and divergence in the mechanisms governing the regulation of sex-biased genes, the complex relationship between the selective forces that drive sex-biased expression and their impact on the process of speciation is still not clearly resolved (Ellegren & Parsch, 2007; Gomes & Civetta, 2015; Harrison et al., 2015; Meiklejohn et al., 2014; Servedio & Boughman, 2017; Servedio & Bürger, 2014).

One strategy to link misregulation to hybrid incompatibility, and hybrid sterility in particular, involves contrasting gene misexpression between sterile and fertile hybrids (Civetta, 2016). Without direct comparison between sterile and fertile hybrids, it is difficult to establish a direct connection between hybrid sterility and gene misregulation (Brekke et al., 2021; Gomes & Civetta, 2014, 2015; Good et al., 2010; Larson et al., 2017; Mack et al., 2016). Consequently, hybrids from reciprocal crosses that exhibit asymmetry in hybrid male sterility – demonstrating Darwin’s corollary to Haldane’s rule – can provide important insights into the molecular mechanisms that cause misexpression and that ultimately are responsible for hybrid dysfunction. Individuals from reciprocal crosses permit a direct comparison of nearly identical F1 hybrid genomes for females and that, for males, only differ in the X-chromosome (Civetta, 2016). While this approach cannot distinguish between sterility-causing genes and the genes they impact, it allows inference of candidate genes likely to be associated with hybrid sterility. Here we investigate the role of regulatory divergence and gene misregulation underlying asymmetric hybrid sterility by comparing gene expression between sterile and fertile hybrids of two *Caenorhabditis* nematodes: *Caenorhabditis remanei* and *C. latens* (Bundus et al., 2018; Dey et al., 2012, 2014).

*C. remanei* and *C. latens* are sister species of nematode roundworms that diverged ∼50 million generations ago (Dey et al., 2012; Félix et al., 2014; Fusca et al., 2025). Like other *Caenorhabditis*, their diploid genomes have five autosome pairs, with males having one hemizygous X-chromosome and females having two X-chromosomes such that X-linked dosage compensation is achieved by reducing the expression of both X-chromosomes in females (Meyer, 2022). Inter-species hybrids can develop to adulthood in both directions of the cross, with hybrid males showing greater hybrid sterility and inviability compared to hybrid females (Haldane’s rule) (Dey et al., 2014) and despite highly conserved outward morphology and undetectable pre-mating or post-mating pre-zygotic reproductive isolation (Dall’Acqua et al., 2025). Reciprocal crosses exhibit an asymmetry in hybrid incompatibility (Darwin’s corollary to Haldane’s rule), with higher hybrid male sterility and inviability with *C. remanei* as the maternal parent (Bundus et al., 2018; Dey et al., 2012, 2014; Dall’Acqua et al., 2025). Previous backcross analysis for this system indicated that the genetic basis for hybrid male sterility is likely simpler than hybrid inviability, involving negative gene interactions between the X-chromosome and autosome(s) (Bundus et al., 2018). Prior research also supports the notion that divergence in sex-biased gene regulation contributes importantly to species-level differences in gene expression between *C. remanei* and *C. latens* (Viswanath et al., 2025). These findings imply that gene regulatory mechanisms have diverged between the two species, which could lead to negative interactions in F1 hybrids, disrupting gene expression and contributing to hybrid incompatibility (Landry et al., 2005a, 2005b; McManus et al., 2010; Landry et al., 2007; Mack et al., 2016; X. C. Li & Fay, 2017). To investigate gene regulatory divergence in asymmetric post-zygotic reproductive isolation between *C. remanei* and *C. latens*, we generated a transcriptomic dataset for both species (*C. remanei* and *C. latens*) and hybrids from both reciprocal cross directions for each sex (males and females) in two tissue types (gonad and soma). By analyzing allele specific expression in hybrids to infer modes of regulatory divergence, we then characterized the evolution of asymmetric misexpression with respect to sex and tissue differences in gene regulation.

## Methods

### Nematode dissection, RNA isolation, and transcriptome sequencing

Following the methods of Viswanath et al. (2025), we dissected male and female nematodes to isolate gonad and soma from F1 hybrid individuals (*C. remanei* VX0003 × *C. latens* VX0088) from both directions of the cross to use for RNA isolation and sequencing. This hybrid data was then integrated with the transcriptome dataset of Viswanath et al. (2025) for comparable sex × tissue sample types from each of the parental species, which were collected contemporaneously. Briefly, we cultured three replicate populations of *C. remanei* (P1), *C. latens* (P2) and F1 hybrids from each reciprocal cross direction (H1 with maternal *C. remanei*, H2 with maternal *C. latens*). We extracted total RNA from ∼400 animals per replicate, including gonad and somatic tissue from male and female adults of wildtype species and hybrids. The F1 male hybrids with *C. remanei* as the maternal parent (H1) had severe gonad defects preventing isolation of gonad tissue for these males, so we additionally collected RNA from whole male worms for *C. remanei*, *C. latens* and hybrid males from reciprocal crosses (H1, H2) to directly compare expression across the different treatments. This collection of samples enabled us to address the effects of tissue type and sex, as well as tissue allometric scaling that can make it challenging to study hybrid misexpression (Assis et al., 2012; Harrison et al., 2015).

All the hybrid samples along with those analyzed in Viswanath et al. (2025) were processed in parallel, being stored in Tri-reagent at -80C for ∼48-72 hours, after which RNA was isolated by Zymo RNA isolation kit (#R2051), treated with Turbo DNase (#AM2238) and RiboLock RNase Inhibitor (#EO0381) and further purified using Zymo RNA Clean and concentrator kit (#R1015) using the Manufacturer’s protocol making a total of 57 samples (3 replicates x 19 treatments), including the pure species samples described in (Viswanath et al., 2025) (Figure 1). These triplicate RNA samples per treatment underwent cDNA library preparation and sequencing at The Centre for Applied Genomics, The Hospital for Sick Children, Toronto, Canada (paired end, read length = 150bp, ∼75 million reads/sample on NovaSeq S4 flowcell).

**Figure 1:**
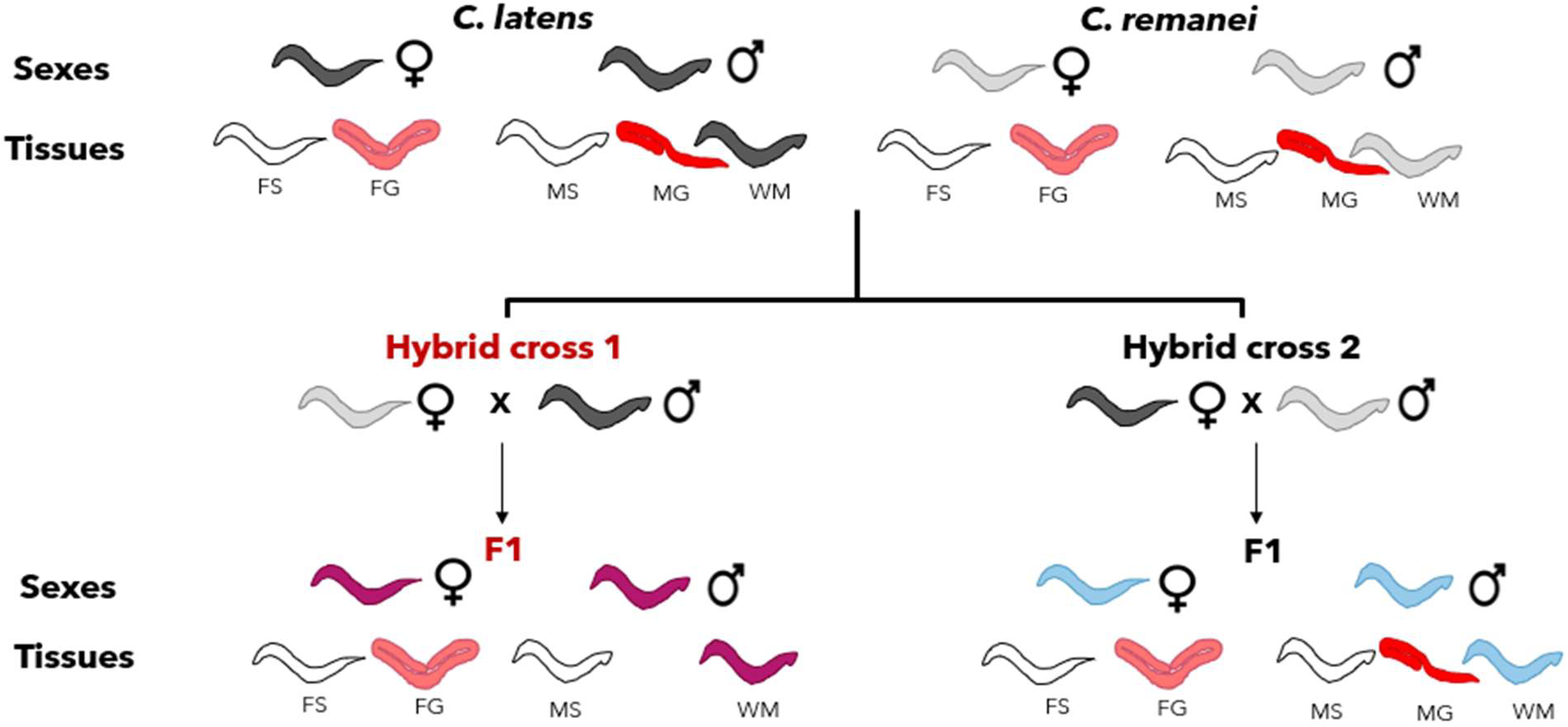
Schematic representation of sample construction. The RNASeq dataset consists of three replicates for each of 19 treatments, comprised of two sexes (females (F) and males (M)) and two tissues per sex (gonad (G) and somatic tissue (S)) derived from each of two wildtype species (*C. latens* (black) and *C. remanei* (grey)), as well as the two reciprocal hybrid classes (H1 cross (pink) and H2 cross (blue)). The exception to this factorial sampling is exclusion of male gonad samples from hybrid cross 1 (H1), due to inability to isolate them reliably. Whole male samples (WM) were also collected for each of the species and hybrid combinations. **Alt text:** Flowchart illustrating tissue sampling (somatic, gonad, whole body) across *C. latens*, *C. remanei*, and reciprocal hybrids, visually highlighting the absence of hybrid cross 1 male gonads.

### Read mapping and allele specific expression analysis

All data processing steps were performed as described previously (Viswanath et al., 2025), including quality trimming of RNAseq reads with Trimmomatic (Bolger et al., 2014)(parameters: LEADING:3 TRAILING:3 SLIDINGWINDOW:4:15 MINLEN:36). Custom gene annotation gff files were generated for both *C. remanei* PX506 (Teterina et al., 2020) and *C. latens* PX534 v2 (Adams et al., 2023) by the Centre for the Analysis of Genome Evolution and Function (CAGEF) with repeat sequences in the genomes being analyzed with RepeatModeler2 using the default parameters (Flynn et al., 2020) and duplicate reads removed with Nubeam-dedup using the default parameters (Dai & Guan, 2020). For more details refer to Viswanath et al., (2025).

The quality verified reads were aligned to the *C. remanei* PX506 (Teterina et al., 2020) and *C. latens* PX534 v2 (Adams et al., 2023) reference genomes using default parameters of STAR (Dobin et al., 2013). The *C. remanei* and *C. latens* samples were mapped to their respective reference genomes while the hybrids were mapped to both *C. remanei* and *C. latens* reference genomes separately. We used CompMap, a Python based competitive read mapping approach (Sanchez-Ramirez & Cutter, 2025; Sánchez-Ramírez & Cutter, 2021) to quantify allele-specific expression (ASE) within each hybrid sample. We then used this quantified ASE to infer genes displaying different regulatory divergence.

Following Sánchez-Ramírez et al. (2021), we used a linear model and a post-hoc Wald type test to test genes for significant *trans* effects (linearHypothesis from the CAR package). We then inferred regulatory divergence based on the statistical criteria defined by McManus et al. (2010) to identify genes displaying divergence due to *cis*-only, *trans*-only and joint *cis*-*trans* effects (Supplementary Figure S1). Genes that exhibited significant expression differences between parental species and hybrid alleles, but lacked significant *trans* effects, were identified as “*cis*-only” divergent. Genes with joint *cis*-*trans* effects were classified as having either synergistic *cis* + *trans* (enhancing) effects, where regulatory divergence of both *cis-* and *trans*-acting factors significantly increases expression of the same allele, or *cis* x *trans* (compensatory) effects, where *cis*-acting regulatory divergence significantly increases expression for the opposite allele as the *trans*-acting regulatory divergence. Genes showing significant expression differences between alleles in hybrids but not between parents were classified as *cis*-*trans* (compensatory). Genes that exhibited no significant expression differences between species and hybrid alleles were classified as “conserved”. All remaining expression comparisons were classified as “ambiguous”. In this way, we inferred regulatory divergence for autosomal genes in both species and X-linked genes in females (Supplementary Figure S1).

For the case of X-linked genes in males, due to the hemizygous nature of the X-chromosome, we were unable to utilize F1 ASE to infer regulatory divergence in the same way as for autosomal loci. However, we used the scheme devised by Sánchez-Ramírez et al. (2021) to confidently identify X-linked genes showing *cis*-only, *trans*-only and *cis*-*trans* (compensatory) divergence as well as conserved genes (Supplementary Figure S2). The prevailing assumption in this scenario is that the *trans*-acting factors affecting X-linked genes primarily come from the autosomes. With this assumption, we classified X-linked genes into different regulatory divergence categories based on their expression with respect to the parental gene expression as given below.

- *X-linked cis-only divergence:* The X-chromosome is inherited from the maternal parent for hybrid males. If X-linked genes exhibit differential male expression between species but equivalent expression for hybrid males and males from the maternal parent species, then it suggests that these genes are primarily influenced by *cis*-only divergence. However, this logic cannot account for the contributions of X-linked and autosomal *trans*-factors, which are similar to those found in the maternal species of hybrids; hence, it is a special case of *cis*-only divergence for X-linked genes in F1 males, represented as *cis*-only*.
- *X-linked trans-only divergence*: These genes display differential gene expression between species, but with X-linked gene expression in hybrids being equivalent to the expression observed for paternal species’ males (X-linked genes in H1 males display expression similar to *C. latens* whereas genes in H2 males display expression similar to *C. remanei*). This subset of genes is expected to show *trans*-only divergence under the assumption that autosomal *C. remanei*-like *trans*-acting factors in H1 and *C. latens*-like *trans*-acting factors in H2 are recessive and not expressed. These X-linked genes are represented as *trans*-only*.
- *X-linked compensatory divergence*: these genes display no differences in expression between species but show higher or lower gene expression than both parents in F1 hybrid males.
- *“Other” X-linked divergence*: Any remaining genes with significant expression divergence were categorized as “other”. These include genes displaying *cis*+*trans* (enhancing), *cis×trans* (compensatory), *trans-*only due to codominant *trans*-regulation divergence and ambiguous expression.

### Differential gene expression analysis and classification of expression in hybrids

To identify gene expression dominance patterns associated with hybrids, we summed up allele-specific counts for each gene for a given hybrid sample. We utilized the DESeq2 package (Love et al. 2014) in R Bioconductor to perform differential gene expression analysis. We then performed a 3-way comparison between *C. remanei*, *C. latens* and Hybrid (H1 or H2) to classify genes in hybrids into different gene expression inheritance categories based on the logic followed by McManus et al. (2010). The genes in hybrids were divided into five categories (Supplementary Figure S3):

- *Dominant expression inheritance*: these genes were significantly differentially expressed between parental species, while in hybrids their expression was identical (i.e., no significant differential expression) to either *C. remanei* (*C. remanei* dominant) or *C. latens* (*C. latens* dominant).
- *Additive expression inheritance:* Genes exhibited significant differences in expression across all three groups, including both parental species and hybrids. However, hybrid expression levels lie between those of the parents.
- *Underdominant expression inheritance:* These genes displayed significant differential expression between parents while also showing significant underexpression in hybrids relative to both parents.
- *Overdominant expression inheritance:* These genes in hybrids demonstrate significant overexpression compared to both parental species, in addition to exhibiting differential expression between the two parental species.
- *Conserved expression inheritance:* These genes displayed no significant expression between all three groups.
- *Ambiguous:* Genes displaying all other expression patterns.

### Data Availability Statement

The mRNA sequencing reads are available under the accession number #GSE328330. All code related to the analyses presented in this study can be found in the following GitHub repository: https://github.com/Athmaja-Viswanath/Gene-regulatory-divergence-underlies-tissue-specific-and-sex-specific-misexpression-in-hybrids

## Results

### Sex specific inheritance and pervasive misexpression of sex-biased genes in hybrids

To discover and investigate the mechanisms of gene regulatory evolution as a potential cause of F1 hybrid incompatibility, we compared transcriptomes between *C. remanei*, *C. latens*, and reciprocally-crossed F1 hybrids for each sex (male, female) and each of two tissue types (gonad, soma). First, by assessing each of the ∼13,000 genes’ expression profiles, we classified genes for each sample type as showing undifferentiated expression of parents and hybrids (“no change”) or into one of five categories for inheritance of expression in hybrids: additive, dominant (*C. latens* dominant or *C. remanei* dominant), and transgressive (underdominant or overdominant).

Our contrasts of expression between males and females revealed clear sex differences in gene expression inheritance in both reciprocal crosses. In the soma tissue samples, males showed ∼1,500 more genes with parent-like “no change” expression profiles than females, indicating that hybrid males more closely resemble parental expression levels despite displaying stronger organismal dysfunction (Chi-square test of independence; H1: p<0.001; H2:p<0.001; n_H1,FS_ = 3044; n_H1,MS_ = 4477; n_H2,FS_ = 3189; n_H2,MS_ = 4777) (Figure 2B, D; Supplementary Figure S4F) (Table 1). Conversely, genes showing dominant expression (*C. remanei* dominant or *C. latens* dominant) are more abundant in hybrid females than males (Chi-square test of independence; H1: p<0.001; H2:p<0.001 for all tissues; n_H1,FS,*Crem*_ = 1582; n_H1,MS,*Crem*_ = 845; n_H2,FS,*Crem*_ = 1444; n_H2,MS,*Crem*_ = 776; n_H1,FS,*Clat*_ = 1147; n_H1,MS,*Clat*_ = 994; n_H2,FS,*Clat*_ = 1304; n_H2,MS,*Clat*_ = 1098) (Figure 2B, D, Supplementary Figure S4F).

**Figure 2:**
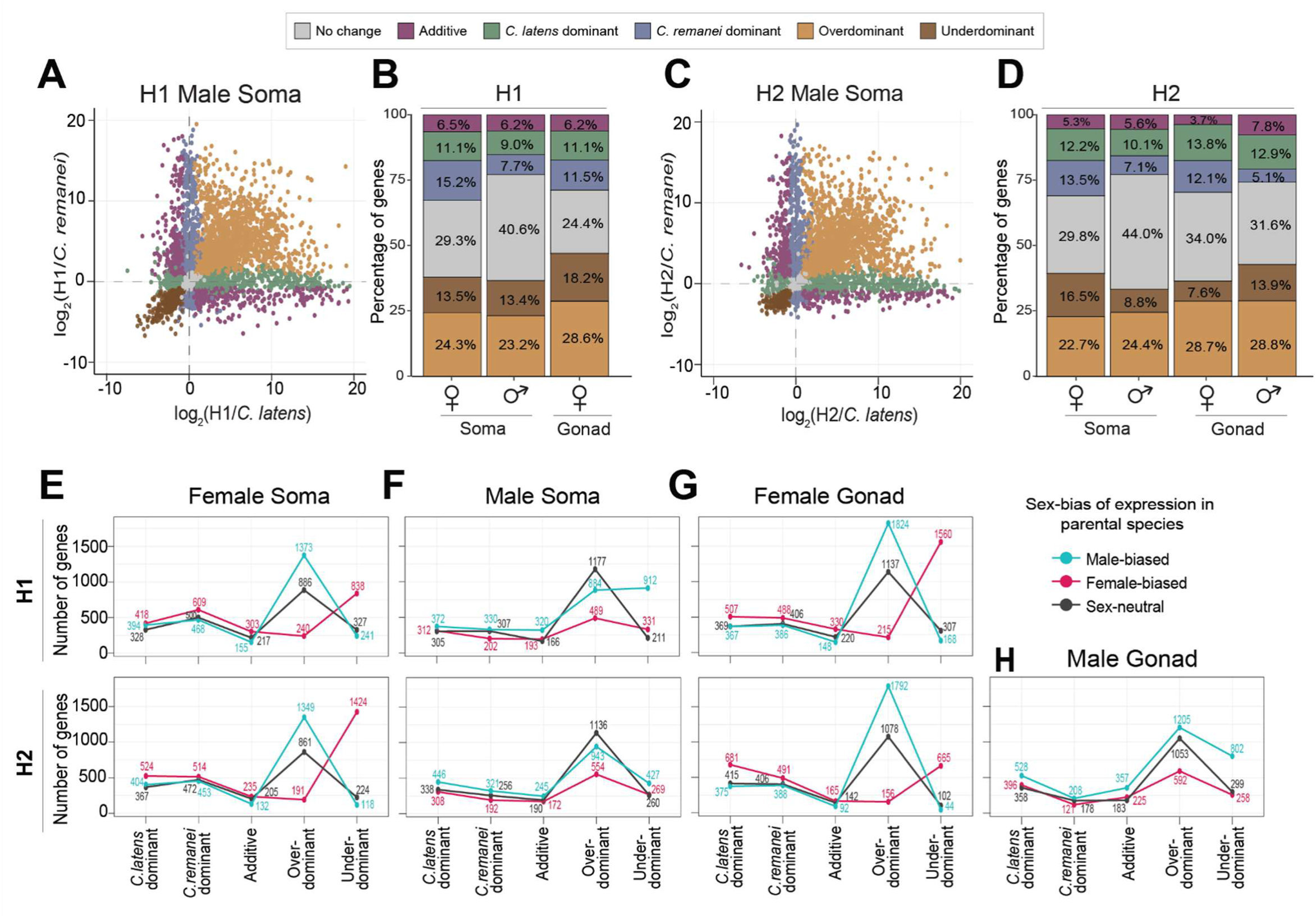
Hybrids reveal extensive transgressive overdominant and underdominant expression in both sexes and for both soma and gonad tissue types. (**A, C**) Scatterplots visualizing different gene expression inheritance categories based on log fold change in soma sample expression for hybrid males relative to each parental species *C. remanei* and *C. latens*. Plots for male gonad expression as well as female gonad and soma expression are shown in Supplementary Figure S4. (**B, D**) Barplots depicting the percentage of genes showing different expression inheritance profiles from each hybrid cross direction for each sex and tissue type. **(E-H)** Transgressive expression in hybrids (overdominant and underdominant) consist mostly of genes with sex-biased expression in the parental species. Plots on the top row correspond to H1 hybrids (*C. remanei* maternal parent) and bottom row corresponds to H2 hybrids (*C. latens* maternal parent). The three lines depict the number of male-biased genes (**blue**), female-biased genes (**pink**) and sex-neutral genes (**black**) observed across inheritance profiles for each sex, tissue type and reciprocal cross. Hybrid females tend to have upregulation of male-biased genes and downregulation of female-biased genes (**E, G**). Hybrid males show misexpression primarily due to sex-neutral and male-biased genes (**F, H**). Genes with the ambiguous expression inheritance category are not shown for clarity. **Alt text:** Multi-panel figure mapping hybrid gene inheritance profiles. Panels A–D use scatterplots and stacked barplots to break down expression categories (like overdominance) by sex and tissue. Panels E–H use line graphs to show that transgressive hybrid expression is heavily driven by parental sex-biased genes.

**Table 1:** Number of genes showing different gene expression inheritance profiles in F1 hybrids.

|  | H1 (*C. remanei* ♀ x *C. latens* ♂) |  |  |  | H2 (*C. latens* ♀x *C. remanei* ♂) |  |  |  |  |
| --- | --- | --- | --- | --- | --- | --- | --- | --- | --- |
| Inheritance Category | Female Soma | Male Soma | Female Gonad | Whole Male | Female Soma | Male Soma | Female Gonad | Male Gonad | Whole male |
| No change | 3044 | 4477 | 2728 | 1989 | 3189 | 4777 | 3619 | 3146 | 1743 |
| *C. latens* dominant | 1147 | 994 | 1247 | 1794 | 1304 | 1098 | 1474 | 1288 | 1768 |
| *C. remanei* dominant | 1582 | 845 | 1286 | 1521 | 1444 | 776 | 1288 | 508 | 1264 |
| Additive | 676 | 686 | 698 | 924 | 572 | 611 | 399 | 755 | 762 |
| Overdominant | 2523 | 2558 | 3203 | 3047 | 2426 | 2649 | 3060 | 2866 | 3453 |
| Underdominant | 1406 | 1479 | 2035 | 2537 | 1766 | 957 | 811 | 1380 | 2689 |
| Ambiguous | 2551 | 2424 | 1870 | 1667 | 2228 | 2596 | 2415 | 2862 | 1800 |
| Total | 12929 | 13463 | 13067 | 13479 | 12929 | 13464 | 13066 | 12805 | 13479 |

Transgressive genes display expression in hybrids beyond the range of both parents (underdominant and overdominant), indicating misexpression in hybrids. Patterns of transgressive misexpression varied between reciprocal crosses: in soma samples for hybrids with the maternal *C. remanei* parent (H1), males exhibited a modest 1% to 5% higher abundance of such genes than females (p=0.0028; n_FS,Underdominant_ = 1406; n_MS,Underdominant_ = 1479; n_FS,Overdominant_ = 2523; n_MS,Overdominant_ = 2558; n_FS,Transgressive_ = 3929; n_MS,Transgressive_ = 4037) (Figure 2B), whereas in H2 soma samples (*C. latens* as maternal parent), hybrid females had substantially more transgressive expression due to ∼1.8-fold more underdominant genes relative to males (p<0.001; n_FS,Overdominant_ = 2426; n_MS,Overdominant_ = 2649; n_FS,Underdominant_ = 1766; n_MS,Underdominant_ = 957; n_FS,Transgressive_ = 4192; n_MS,Transgressive_ = 3606) (Figure 2D) (Table 1). This elevated female-biased misexpression was unexpected, given that H2 males display only slightly stronger hybrid dysfunction than their hybrid sisters, but the pattern parallels observations in *C. nigoni - C. briggsae* hybrids (Sánchez-Ramírez et al., 2021).

In the case of gonad tissue samples of hybrids, we compared male versus female gonad expression only in the H2 cross because H1 male gonad tissues were too underdeveloped to isolate and analyze reliably. Gonadal expression in H2 hybrids further revealed strong sex-specific regulatory disruption, with hybrid males displaying twice the number of additive genes (n_FG_ = 399; n_MG_ = 775) and ∼1.7 fold higher abundance of underdominant genes as hybrid females (p<0.001; n_FG,Overdominant_ = 3060; n_MG,Overdominant_ = 2866; n_FG,Underdominant_ = 811; n_MG,Underdominant_ = 1380; n_FG, Transgressive_ = 3871; n_MG, Transgressive_ = 4246) (Figure 2D; Supplementary Figure S4G). Conversely, H2 hybrid females exhibit significantly more genes with parent-like expression than their brothers (p<0.001; n_FG_ = 3619; n_MG_= 3146) as well as more genes displaying simple expression dominance (p<0.001; n_FG,*Crem*_ = 1288; n_MG,*Crem*_ = 508; n_FG,*Clat*_ = 1474; n_MG,*Clat*_ = 1288) (Figure 2D; Supplementary Figure S4G) (Table 1). Finally, H2 male gonads express ∼20% more transgressive genes than seen for H2 male soma (p<0.001; n_MS_ = 3606; n_MG_ = 4246) (Figure 2D; Supplementary Figure S4I), consistent with higher regulatory sensitivity in reproductive tissues (Cutter, 2023b; Cutter & Bundus, 2020; de Zwaan et al., 2022).

Female hybrids derived from *C. remanei* and *C. latens* more rarely experience sterility and inviability than do hybrid males (Dey et al., 2014). To gain insights into the gene regulatory mechanisms that might underlie sex-biased hybrid incompatibility, we examined sex-biased misexpression in hybrids with reference to sets of genes having sex-biased expression that we previously defined for the parental species *C. remanei* and *C. latens* (Viswanath et al., 2025). We discovered that a majority of the transgressive genes in both males and females were sex-biased in parental expression (Figure 2 E-H). Specifically, for hybrid females, more than half of overdominant genes showed male-biased expression in the parental species, indicating that genes that typically show male-biased expression tend to get upregulated in hybrid females (Figure 2E, G; blue line). Conversely, ∼60-80% of underdominant genes display female-biased expression in the parents, indicating that genes that typically are female-biased get downregulated in hybrid females (Figure 2E, G; pink line). These patterns remain consistent for female gonad and soma, as well as reciprocal crosses, and suggest that transgressive expression in hybrid females leads to masculinization of the female transcriptome by preferentially upregulating male-biased genes while simultaneously downregulating female-biased genes.

Hybrid males show transgressive misexpression of those genes that ordinarily would be male-biased and sex-neutral in the parental species (Figure 2F, H). Genes with underdominant expression in hybrid males, in particular, are highly enriched for those genes that typically have male-biased expression (n_male-biased_= 802; n_sex-neutral_ = 299; n_female-biased_ = 258), suggesting that misexpression in hybrid males is due to suppressed expression (Figure2 F, H; blue line; p_H1,MS_ <0.001; p_H2,MS_ = 0.01; p_H2,MG_ <0.001). Such misexpression of male-biased genes in hybrids also has been documented in other taxonomic groups including flies, mice, and other nematodes (Gomes & Civetta, 2015; Mack et al., 2016; Sánchez-Ramírez et al., 2021).

Despite the pervasive misexpression of sex-biased genes in both sexes, organismal hybrid incompatibility is more severe in males than females. The misexpression of male-biased genes in both sexes further suggests that male-biased gene networks may be especially prone to both misexpression and dysfunctional organismal outputs, whereas female-biased gene networks tend to be more resilient to perturbations. Moreover, male-biased gene regulatory networks may be “fragile” due to faster molecular evolution of male-biased genes or due to unique selective pressures driving the evolution of sex-biased gene expression (Wu & Davis 1993; Delph & Demuth 2016; Grath & Parsch 2016; Cutter & Bundus 2020; Cutter 2023).

### Parent-of-origin asymmetry in hybrid misexpression biased toward *C. remanei*-maternal (H1) hybrids

Hybrid male sterility is more common in the H1 cross (*C. remanei* as the maternal parent) compared to H2 cross (with *C. latens* as the maternal parent) (Bundus et al., 2018; Dey et al., 2012, 2014), leading us to examine gene expression patterns of hybrids for sensitivity to cross-direction for a given sex and tissue (female gonad, female soma, male soma). Across all three sex × tissue combinations that we could analyze, H1 hybrids showed fewer genes with expression equivalent to the parentals than did H2 hybrids especially in the female gonad (Supplementary Figure S5; p_FS_ = 0.78; p_MS_ = 0.55; p_FG_ < 0.001; p_WM_ <0.001). This cross-direction difference is indicative of overall greater disruption of expression in H1 hybrids and was especially pronounced in the female gonad with ∼890 fewer genes in H1 than H2 (p_FG_ < 0.001; female soma n_H1_=3044 or ∼29%; n_H2_=3189 or ∼30%; male soma n_H1_ = 4477 or ∼40.5%; n_H2_ = 4777 or ∼44%; female gonad n_H1_=2728 or ∼24%; n_H2_=3619 or ∼34%) (Supplementary Figure S5). Moreover, H1 hybrids also exhibited more -genes with additive expression profiles than H2 hybrids (p_FS_ = 0.002; p_MS_ = 0.03; p_FG_ < 0.001; p_WM_ <0.001; female soma n_H1_ = 676 or ∼6.5%; n_H2_ = 572 or ∼5%, Supplementary Figure S5A; male soma n_H1_ = 686 or ∼6%; n_H2_ = 611 or ∼5.6%, Supplementary Figure S5B; female gonad, n_H1_ = 698 or ∼6%; n_H2_ = 399 or ∼4%,Supplemenatry Figure S5C). Additivity results in intermediate gene expression levels that nonetheless deviate from the expression profiles of both parental species and thus could contribute to dysfunction of genetic pathways. Finally, and more critically, H1 hybrids showed a substantially higher incidence of transgressive genes than H2 in the male soma (p_MS_ < 0.001; n_H1_=4037 or ∼36%; n_H2_=3606 or ∼33%) and female gonad (p_FG_ < 0.001; n_H1_=5238 or ∼47%; n_H2_=3871 or ∼36%) (Supplementary Figure S5B, C). Specifically, genes with underdominant expression were ∼1.5 fold more common in H1 than H2, contributing to the elevated transgressive expression of H1 (Supplementary Figure S5B, C). Overall, these results indicate that the H1 hybrid cross, which experiences a higher incidence of hybrid male sterility with *C. remanei* as the maternal parent, has more profound misexpression than hybrids from the reciprocal H2 cross, regardless of sex or tissue type.

We next aimed to determine whether the same genes were misexpressed in both reciprocal hybrid crosses to understand whether or not the underlying regulatory mechanisms were unique to each hybrid cross. We observed a significant overlap of genes in most expression dominance categories between hybrid crosses (Figure 3A-C; Chi-square test of independence, p <0.001 for all tissues). In hybrid females, however, of the transgressive genes that showed different expression inheritance in each hybrid cross, ∼48-58% of them displayed transgressive expression in only one of the two hybrid crosses while displaying parent-like expression in the alternative cross (n_FS,H1_ = 258/444, ∼58%; n_FS,H2_ = 252/466, ∼54%; n_FG,H1_ = 448/823, ∼54%; n_FG,H2_ = 46/95, ∼48%) (Figure 3A, C). This pattern of transgressive expression being unique to genes in one cross-direction was even more extreme in male soma (n_MS,H1_ = 951/1118, ∼85%; n_MS,H2_ = 642/818, ∼78%) (Figure 3B). These observations clearly indicate that many distinct genes are misexpressed in each hybrid cross. A notable genetic distinction between the H1 and H2 males is that they inherit their X-chromosome and mitochondrial genome from *C. remanei* versus *C. latens*, respectively, which could underlie their different patterns of misexpression. Although we did not observe any obvious chromosomal enrichment of transgressive genes (Supplementary figure S6), the difference in X-chromosome origin might nonetheless indirectly influence the unique misexpression of genes in H1 and H2 male hybrids, potentially underlying the asymmetric hybrid incompatibility.

**Figure 3:**
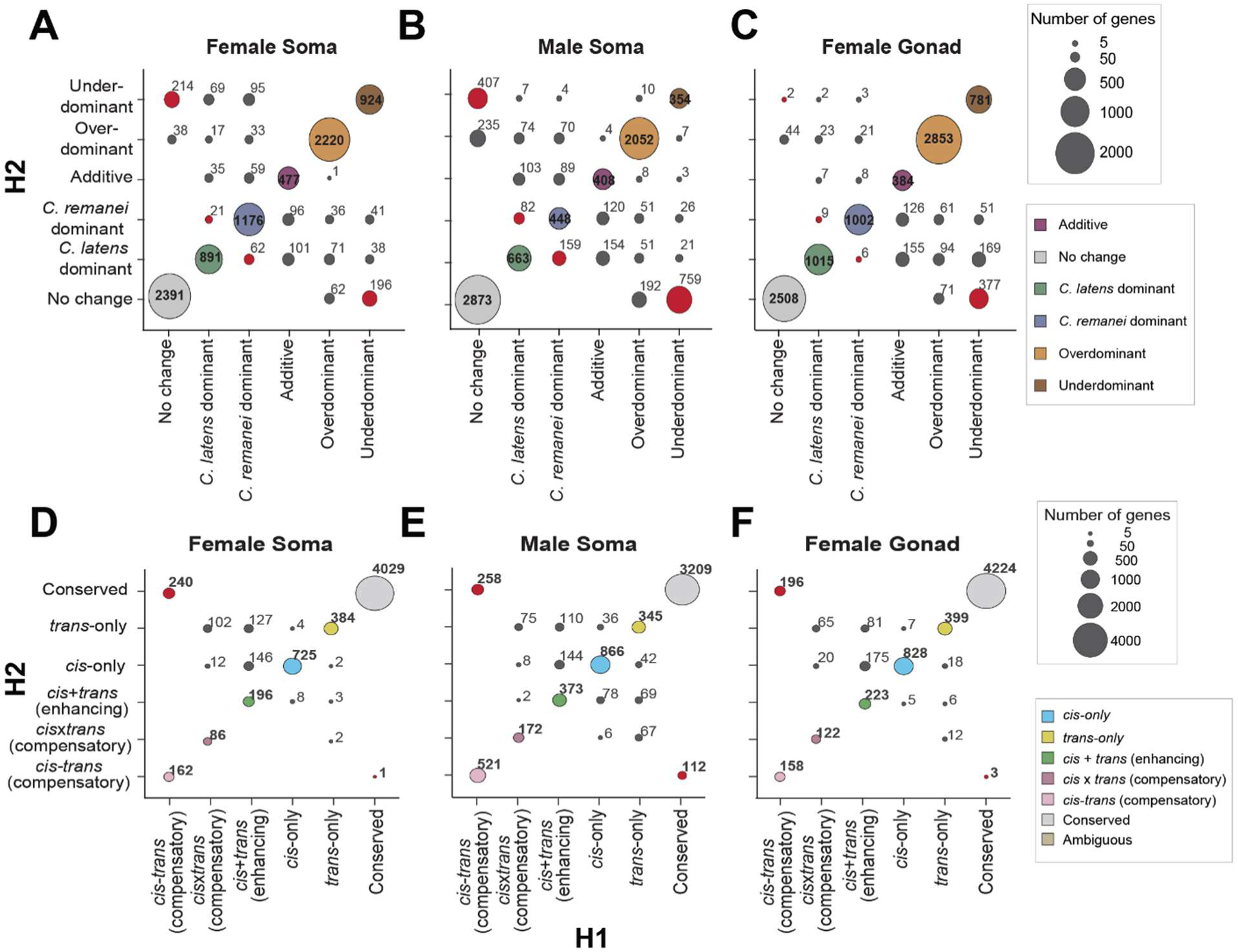
Despite higher overlap in genes across reciprocal crosses, H1 and H2 display distinct patterns for expression inheritance and regulatory divergence. (**A, B, C**) Bubble plots contrasting the genes in each expression inheritance category across reciprocal crosses with area of the bubble proportional to the number of genes. The bubbles on the diagonal represent the number of genes overlapping between H1 (x-axis) and H2 (y-axis) and red bubbles highlight the number of genes showing underdominant expression exclusively in one hybrid cross. Females show a higher level of overlap in genes than males across reciprocal crosses (**A, C**, bubbles on the diagonal). Males have higher number of underdominant genes unique to each hybrid cross (**B**, red bubbles). (**D, E, F**) Bubble plots contrasting the genes and their regulatory divergence in H1 (x-axis) and H2 (y-axis). The bubbles on the diagonals represent the number of genes overlapping between H1 (x-axis) and H2 (y-axis) and red bubbles highlight the number of genes showing *cis*-*trans* compensatory divergence exclusively in one hybrid cross. More than 50% of genes showing *cis*-*trans* compensatory regulation in H1 females were conserved in H2 females. Only autosomal genes are represented for clarity as the X-linked genes were classified with a slightly different criteria due to their hemizygous nature in males (see methods; Supplementary Figure S2). Both autosomal and X-linked genes display similar patterns for male soma, however, in case of X-linked genes in female soma and gonad, no deviation is observed from the diagonal indicating hybrid-cross specific misregulation predominantly of autosomal genes in females (Supplementary Figure S11). **Alt text:** Multi-panel bubble plot matrix mapping concordance between H1 and H2 reciprocal crosses. Panels A–C compare expression inheritance categories, while D–F compare regulatory divergence mechanisms. Off-diagonal circles (highlighted in red) visually isolate genes with hybrid-cross-specific misregulation across female soma, male soma, and female gonad tissues.

### *cis*-only divergence is more prevalent than *trans*-only divergence

We used allele-specific expression analysis with the F1 hybrids to classify genes as having evolved distinct categories of regulatory divergence: *cis*-only, *trans*-only, *cis*-*trans* compensatory, *cis*+*trans* (enhancing) and *cis×trans* (compensatory) (Supplementary figure S1). In hybrids, consistently more autosomal genes were affected by divergence in *cis* than *trans*, whether considering all genes showing regulatory divergence or just those genes with *cis*-only or *trans*-only divergence (Table 2). Specifically, expression profiles in both males and females in reciprocal crosses indicated almost twice as many genes having experienced *cis*-only divergence (∼8-16% of genes) compared to *trans*-only divergence (∼5-7% of genes) (Table 2; Figure 4B, D). Within the H1 cross, *cis*-only divergence was especially abundant in female soma relative to male soma (p_Soma_ < 0.001; n_FS_ = 1700 or ∼16%; n_MS_ = 1104 or ∼10%) (Figure 4A, B; Supplementary Figure S7F). Conversely, the H2 cross revealed a higher incidence of genes with *cis*-only divergence in males than females for both soma and gonad tissues (p_Soma_ < 0.001; n_FS, *cis*-only_ = 891 or ∼8%; n_MS, *cis-*only_ = 1132 or ∼10%; p_Gonad_ < 0.001; n_FG, *cis*-only_ = 1093 or ∼10%; n_MG, *cis-*only_ = 1223 or ∼12%) (Figure 4C, D Supplementary Figure S7F) (Table 2). Genes that exhibited *trans*-only divergence, however, were similar in abundance across sexes and hybrid crosses, comprising <10% of all the genes analyzed (n_H1,FS, *trans*-only_ = 659 or ∼6%; n _H1,MS, *trans*-only_ = 550 or ∼5%; n_H2,FS, *trans*-only_ = 622 or ∼6%; n _H2,MS, *trans*-only_ = 584 or ∼5.4%; n_H2, FG, *trans*-only_ = 554 or ∼5%; n _H2, MG, *trans*-only_ = 530 or ∼5%) (Table 2). These general patterns also held when restricting the analysis to X-linked loci (Table 3; Supplementary Figure S8). These patterns for *C. remanei* and *C. latens* are consistent with the classic expectation that *cis*-regulatory elements will disproportionately accumulate differences between species over the long term, as a result of selective filtering of their more limited pleiotropic mutational effects (Bell et al., 2013; Mack et al., 2016; Wittkopp et al., 2008; Carroll, 2005; Stern & Orgogozo, 2009). However, previous work with whole animal transcriptomic data in *C. briggsae* and *C. nigoni* nematodes identified a larger contribution of *cis*-*trans* compensatory changes in hybrid incompatibility (Sánchez-Ramírez et al., 2021; Viswanath & Cutter, 2023), a point that we return to in the Discussion.

**Figure 4:**
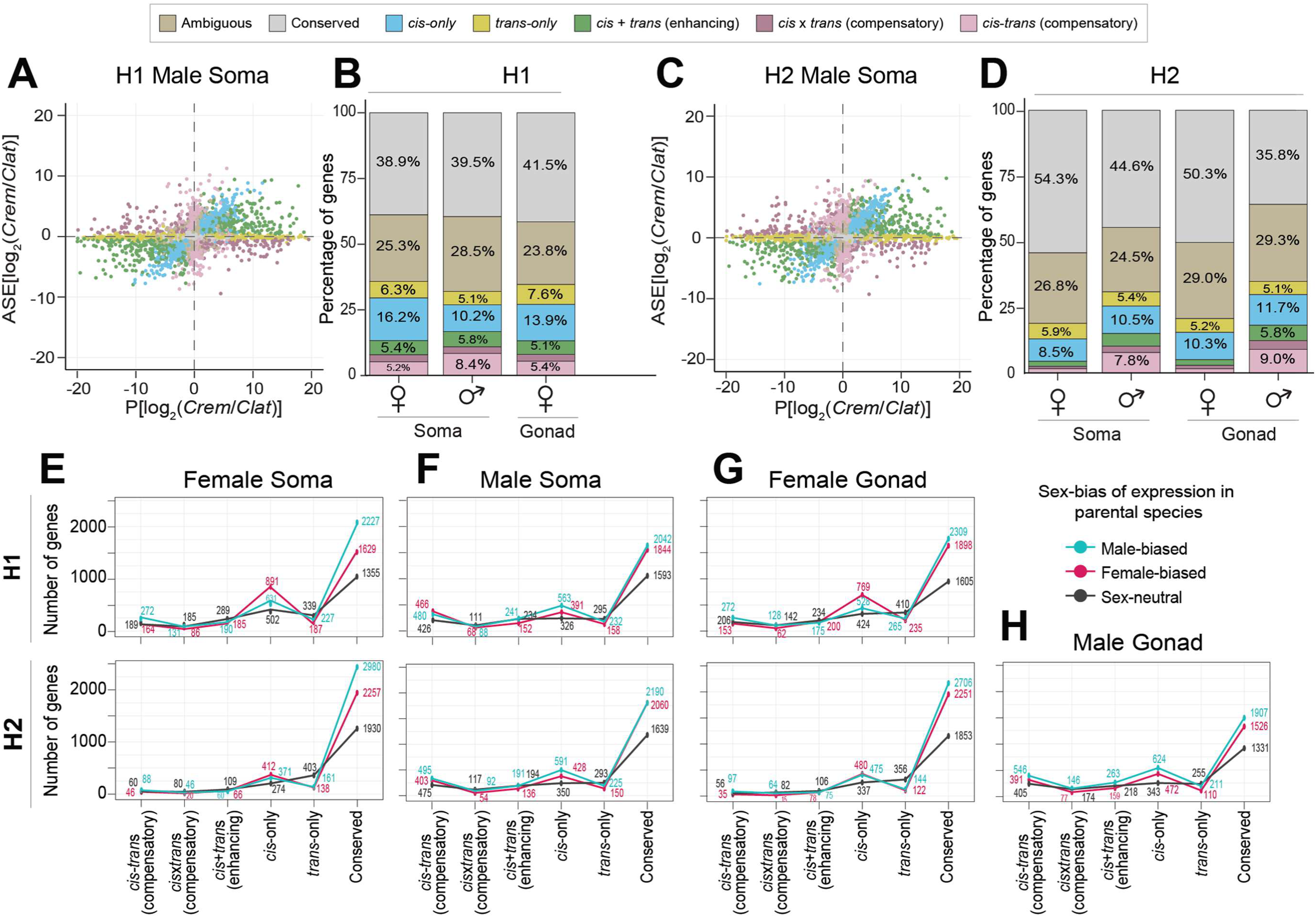
Gene misregulation differs between sexes and reciprocal hybrid crosses. (**A, C**) Classification of regulatory divergence using allele specific expression analysis. Gene expression between the species *C. remanei* (*Crem*) and *C. latens* (*Clat*) (x-axis) is compared to the expression difference between alleles within the hybrids (y-axis). Genes with conserved regulation across species are depicted in grey. Such genes show no expression difference between species as well as no difference between alleles within hybrids and fall on the origin (0,0). Scatterplots depicting regulatory divergence for genes expressed in male gonad, female soma and gonad are depicted in Supplementary Figure S7. (**B, D**) Barplots depicting the percentage of genes showing different regulatory divergence categories detected in each reciprocal cross H1 (maternal *C. remanei* hybrids) (**B**) and H2 (maternal *C. latens* hybrids) (**D**) for autosomal genes. More than ∼38% of total genes in all samples displayed allele-specific expression profiles consistent with regulatory conservation. H2 males displayed an overall higher number of genes showing regulatory divergence (predominantly *cis*-only and *cis*-*trans* compensatory divergence) than females (n_MS_ = 3341; n_FS_ = 1979; n_MG_ = 3655; n_FG_ = 2184). This pattern also broadly holds true for X-linked genes (Supplementary Figure S8). (**E-H**) Sex-biased genes display conserved regulation or *cis*-only regulatory divergence. *cis*-only divergence disproportionately influences female-biased genes in hybrid females (**E, G**) and male-biased genes in hybrid males (**F, H**). In contrast, *trans*-only divergence tends to affect sex-neutral genes in both sexes. Genes with ambiguous expression categorization not shown for clarity. Plots on the top row correspond to H1 hybrids (*C. remanei* maternal parent) and bottom row corresponds to H2 hybrids (*C. latens* maternal parent). While this line plot represents autosomal genes, it is important to note that *cis*-*trans* compensatory divergence also plays a role in the misexpression of sex-biased genes, particularly in males with respect to X-linked genes (Supplementary Figure S9). **Alt text:** Multi-panel figure analyzing hybrid regulatory divergence. Panels A–D use scatterplots and stacked barplots to quantify *cis* and *trans* regulatory categories by sex and tissue. Panels E–H use line graphs to show how parental sex-biased genes track with specific regulatory mechanisms like cis-only divergence.

**Table 2:**
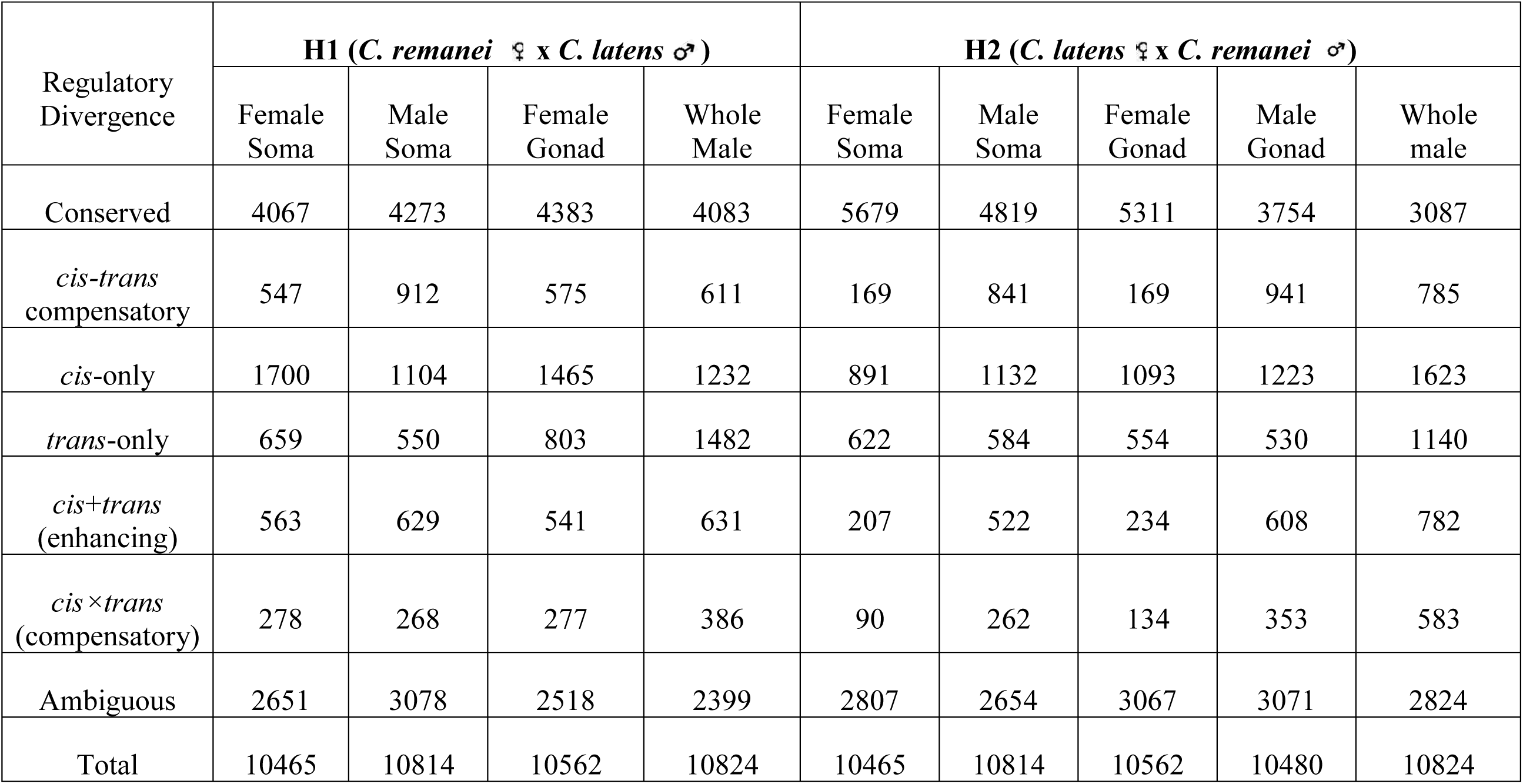
Number of autosomal genes showing regulatory divergence in F1 hybrids.

**Table 3:** Number of X-linked genes showing regulatory divergence in F1 hybrids.

| Regulatory Divergence | H1 (*C. remanei* ♀ x *C. latens* ♂) |  |  |  | H2 (*C. latens* ♀ x *C. remanei* ♂) |  |  |  |  |
| --- | --- | --- | --- | --- | --- | --- | --- | --- | --- |
|  | Female Soma | Male Soma | Female Gonad | Whole Male | Female Soma | Male Soma | Female Gonad | Male Gonad | Whole male |
| Conserved | 1175 | 1224 | 1460 | 440 | 1531 | 1097 | 1541 | 1023 | 314 |
| *cis-trans* compensatory | 77 | 459 | 54 | 839 | 24 | 527 | 18 | 405 | 935 |
| *cis*-only | 326 | 177* | 254 | 288* | 162 | 239* | 194 | 222* | 369* |
| *trans*-only | 95 | 136* | 108 | 292* | 81 | 87* | 70 | 47* | 176* |
| Other | 771 | 628 | 604 | 767 | 646 | 674 | 657 | 629 | 832 |
| Total | 2444 | 2624 | 2480 | 2626 | 2444 | 2624 | 2480 | 2326 | 2626 |

### Compensatory *cis*-*trans* changes underlie differences between males and females

To further explore the evolution of gene regulation between *C. remanei* and *C. latens*, we assessed the relative impact of sex-specific divergent *cis-* and *trans-*acting regulators on F1 hybrids through allele specific expression analysis. First, we found that *cis*-only divergence disproportionately revealed itself in hybrid females for genes that are typically female-biased in the parental species (Chi-square tests, p < 0.001 for all somatic and gonadal tissues across both crosses; Figure 4E, G). Complementarily, hybrid males revealed *cis*-only divergence for genes that typically have male-biased expression in the parental species (Figure 4F, H). In contrast, *trans*-only divergence was associated with sex-neutral genes in both males and females (Figure 4E-H). These observations indicate that *cis-* versus *trans*-acting regulatory divergence also play distinct roles in sex-biased gene regulation to contribute to species differences in gene expression.

Next, we found that gene expression of the sexes differed in the incidence of genes with detectable joint effects of both *cis*-acting and *trans*-acting regulatory divergence. Genes exhibiting *cis-trans* compensatory regulatory changes, in particular, were ∼1.5 times more common in males than females in H1 soma (n_FS_ = 547 or ∼5%; n_MS_ = 912 or ∼8%) and ∼7 times as abundant in H2 males as females for autosomal genes (n_FS_ = 169 or ∼1%; n_MS_ = 841 or ∼7%) (Figure 4B, D; Supplementary Figure S7F) (Table 2). Among the gonad genes showing any form of regulatory divergence, males expressed ∼5.5-fold more genes with *cis-trans* compensatory divergence than the female transcriptome, consistent with H2 soma patterns (n_FG, Compensatory_ = 169 or ∼1%; n_MG, Compensatory_ = 941 or ∼9%) (Figure 4D, Supplementary Figure S7 G). Furthermore, male gonads also showed a higher abundance of genes in all other regulatory divergence categories than females, consistent with males showing overall more extensive regulatory divergence than females (Figure 4D, Supplementary Figure S7G; Table 2). In the case of autosomal genes with conserved regulation for gonad tissue samples, by contrast, female H2 hybrids had ∼1.4-fold more than males, similar to soma tissue (n_FG_ = 5311 or ∼50%; n_MG_ = 3754 or ∼36%) (Figure 4D, Supplementary Figure S7G) (Table 2). The disproportionate abundance of compensating *cis-trans* differences evident in males compared to females for both hybrid crosses and in both tissues shows that *cis*-*trans* compensatory evolution is likely driving key sex differences in gene regulatory controls.

Differences in regulatory divergence patterns between males and females could be attributed to the varying effects of X-linked genes on each sex, as male (XO) and female (XX) *Caenorhabditis* differ in their number of X-chromosome copies. Due to the hemizygous nature of the X-chromosome in males, we used a modified classification scheme based on gene expression inheritance following Sánchez-Ramírez et al. (2021) (see Methods) to categorize X-linked genes as *cis*-only*, *trans*-only*, and *cis*-*trans* compensatory divergence, as well as conserved and “other” [*cis*+*trans* (enhancing), *cis×trans* (compensatory), ambiguous]. The use of modified criteria for classifying X-linked genes is indicated by the asterisks (*), although this re-categorization is conceptually consistent with the categories defined in females. In line with patterns observed for autosomes, X-chromosomes were 2-to-3-fold enriched for genes displaying *cis*-*trans* compensatory divergence from male transcriptomes versus female transcriptomes showing 1.6-fold to 2-fold underrepresentation of such genes on the X-chromosome (Fisher exact test, p < 0.05) (Figure 5). In addition, genes displaying conserved regulation show a ∼1.4**-**fold to 2-fold enrichment on the X-chromosomes for females and modest increase in abundance on the X-chromosome for males (Figure 5). These patterns implicate an especially large role of the X-chromosome in *cis*-*trans* compensatory regulatory divergence in males, as also seen in hybrids of *C. briggsae* × *C. nigoni* (Sánchez-Ramírez et al., 2021).

**Figure 5:**
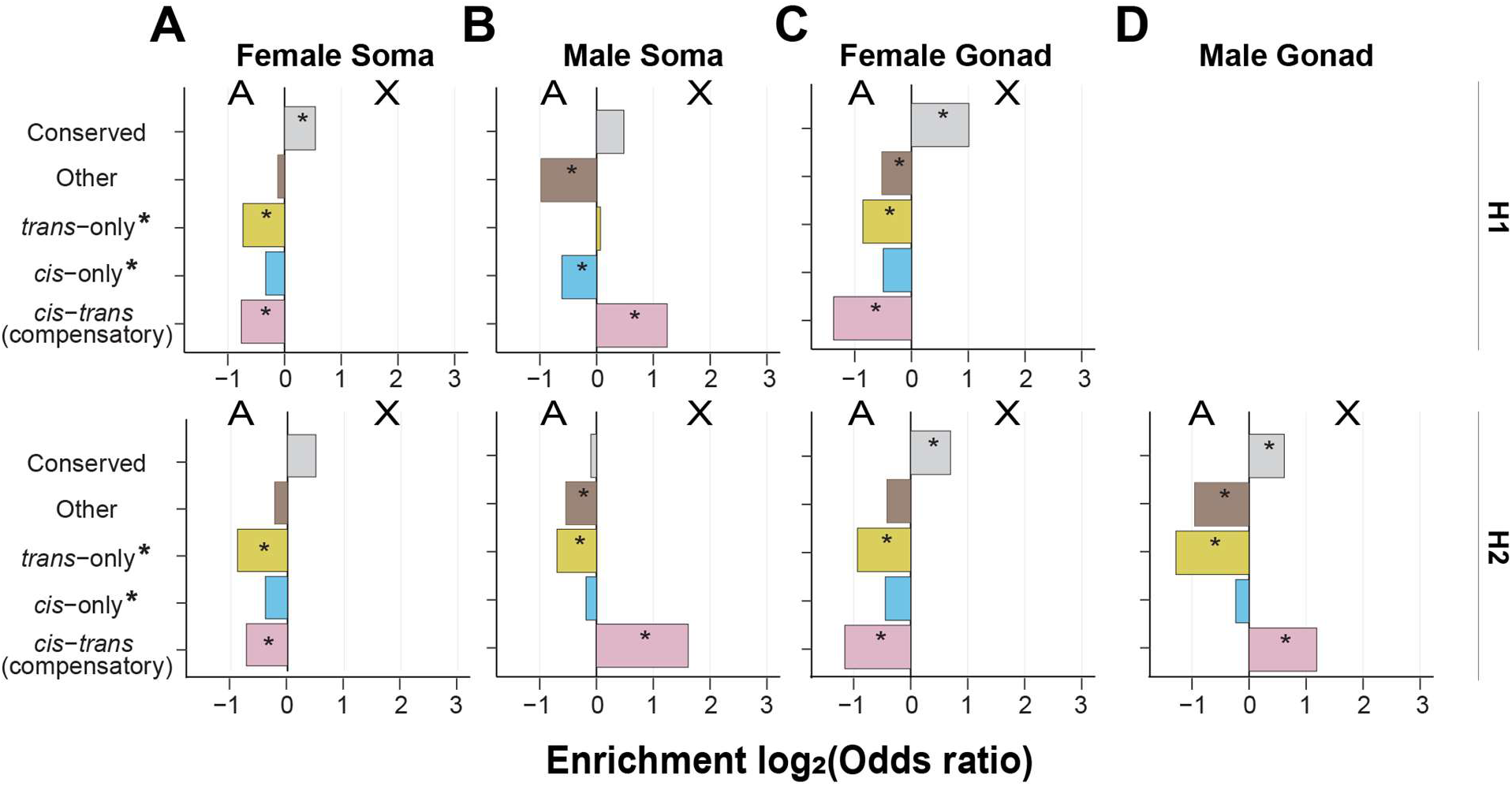
*cis*-*trans* compensatory changes are enriched on the male X-chromosome. Enrichment of genes linked to autosomes versus X-chromosomes for different categories of regulatory divergence inferred from soma (**A, B**) and gonad (**C, D**) tissue types of females (**A, C**) and males (**B, D**) for H1 hybrids (top row, maternal *C. remanei* hybrids) and H2 hybrids (bottom row, maternal *C. latens* hybrids). H1 hybrids have *C. remanei* maternal parents; H2 hybrids have *C. latens* maternal parents. Positive values of the log_2_(odds ratio) denote enrichment for the X-chromosome and negative values indicate enrichment for autosomes. The X-chromosome of males exhibited an enrichment of genes displaying *cis*-*trans* compensatory regulatory divergence, whereas such genes were underrepresented on the X-chromosomes of females. Regulatory categories defined according to the restricted criteria used for X-chromosomes in males (Supplementary Figure S2). **Alt text:** Grid of horizontal bar charts displaying chromosomal enrichment odds ratios by regulatory category. Rows contrast H1 and H2 crosses across female/male soma and gonad tissues, visually highlighting X-chromosome enrichment for male *cis-trans* compensatory changes and noting the absence of H1 male gonad data.

### Distinct gene regulatory mechanisms underlie misexpression in hybrids from reciprocal crosses

We next compared reciprocal crosses in terms of which genes were inferred to correspond to a given regulatory divergence category to understand how gene misregulation might underlie the asymmetry in hybrid incompatibility. To do so, we tested whether regulatory divergence affected the same or distinct sets of genes in H1 and H2. In the male soma, each regulatory divergence category showed an overlap of >50% of genes between H1 and H2. For genes inferred to have *cis*-*trans* compensatory regulatory changes, in particular, we found similar gene abundances and ∼67%-82% overlap in gene identity between H1 and H2 for the male soma samples (n_H1_ = 521 of 779, 67%; n_H2_ = 521 of 633, 82%; Figure 3E).

Genes with unique and cross-direction-specific *cis*-*trans* compensatory effects, however, were nearly twice as common for H1 male soma than for H2 male soma (n_H1_ = 258; n_H2_ = 112; Figure 3E, red bubbles). These hybrid-cross specific genes account for ∼33% and ∼18% of all the genes showing *cis*-*trans* compensatory divergence in H1 and H2, respectively, and display conserved regulation in the corresponding reciprocal cross (Figure 3E). Genes with cross-specific effects (Figure 3E, red circles) may disproportionately influence gene expression globally, resulting in distinct patterns of gene misregulation in reciprocal crosses. If so, male hybrids from both cross directions may largely share the same mechanism of expression disruption despite their contrasting severity of organismal hybrid incompatibility, possibly involving relatively few genes unique to each hybrid cross. Alternatively, the substantial overlap in genes inferred to have the same regulatory divergence category for H1 and H2 male soma might simply be a byproduct of tissue similarity, as hybrid male somatic bodies show less obvious and severe dysfunction compared to gonads (Dall’Acqua et al., 2025). Unfortunately, we were unable to definitively discern this aspect of asymmetry due to the lack of reliable hybrid gonad data for H1 males.

In females, more than ∼55% of the autosomal genes exhibiting *cis*-*trans* compensatory regulatory divergence were unique to H1 and did not overlap the gene inferences from H2 (Figure 3D, F), despite the nuclear genome being identical for both classes of F1 female. When we restricted our analysis to X-linked loci, however, we found that all the genes overlapped in identity, suggesting that hybrid-cross specific misregulation predominantly involves autosomal genes in H1 females (with *C. remanei* as the maternal parent) (Supplementary Figure S11). Additionally, we noticed that more autosomal genes change from *cis*-only divergence in one hybrid cross to *trans*-only divergence in the reciprocal cross for male soma compared to female soma (Figure 3D-F), even though more than half of the genes exhibiting *cis*-only and *trans*-only divergence overlapped between H1 and H2 in both sexes. These distinctive gene misregulation patterns support the notion that uniparentally inherited factors influence gene regulatory divergence inferences in both sexes, with parent-of-origin effects potentially involving X-chromosomes, mitochondria, histone modifications, or small RNA pathways.

### Distinct gene regulatory divergence underlying sex-dependent transgressive expression

Our next objective aimed to identify the gene regulatory changes that led to transgressive expression in hybrids that exceeded parental expression levels, in order to gain insights into the role of *cis-* and *trans-*acting mechanisms in hybrid misexpression. We found different gene regulatory mechanisms to underlie transgressive expression in male versus female hybrids, mirroring our previous observation of distinct genes with transgressive expression in each sex. In males, compensatory *cis-trans* regulatory divergence underlies a higher proportion of transgressive genes than other regulatory divergence categories (Figure 6). In females, like males, compensatory *cis-trans* contributed importantly to transgressive hybrid expression profiles, but substantial transgressive expression also was due to other regulatory divergence categories like *cis*-only and *tran*s-only (Chi-square tests; H1, MS*χ*^2^ = 1666.5; H2, MS*χ*^2^= 1848.6; H2, MG*χ*^2^ = 1856.4; p < 0.001 for all; Figure 6). The difference in relative importance of gene regulatory divergence categories that underlie transgressive hybrid expression detected in males versus females further indicates that *cis-* and *trans*-acting regulators play distinct roles in male and female genetic networks.

**Figure 6:**
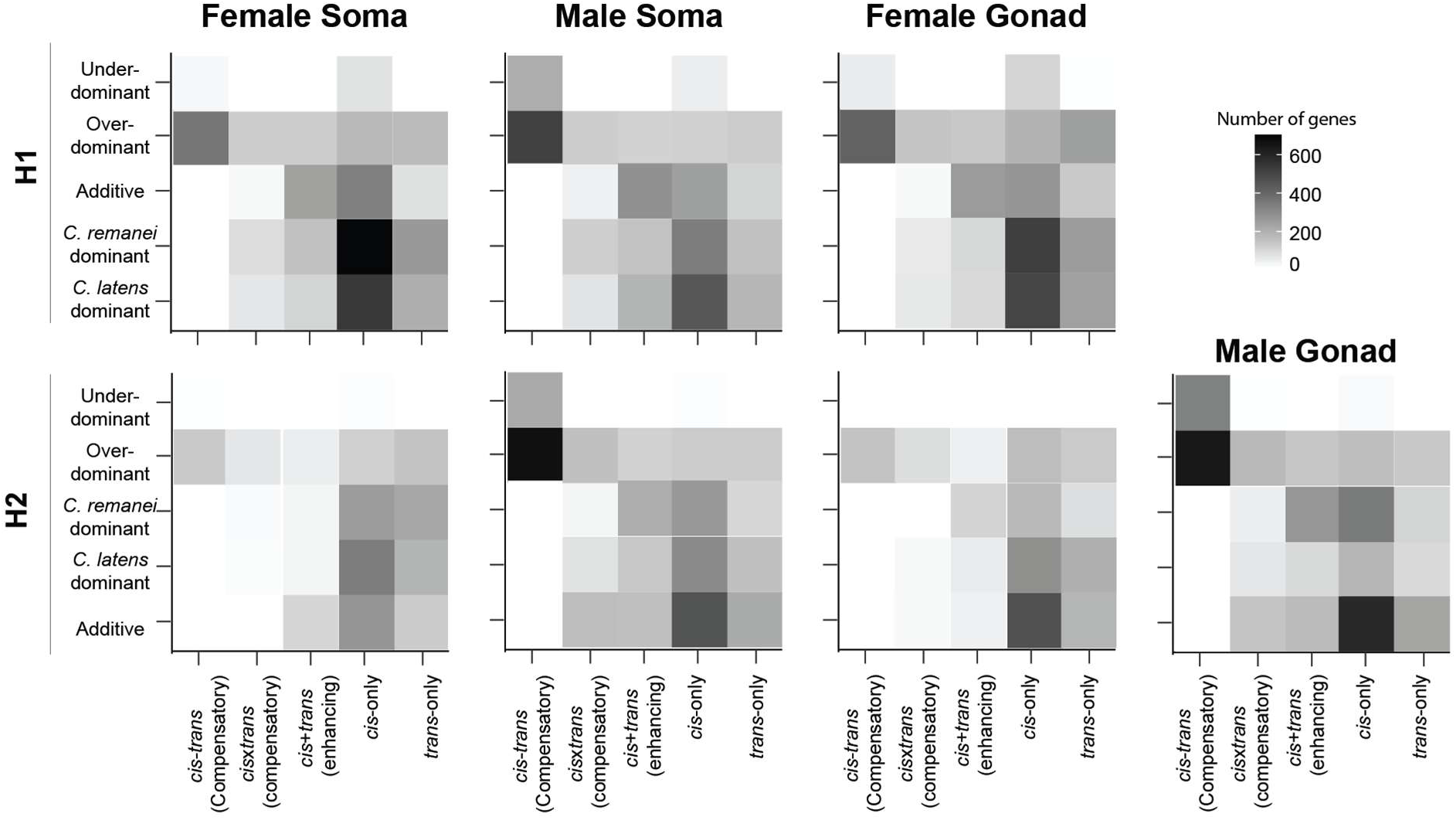
*cis*- and *trans*- changes underlie higher proportion of transgressive genes in both sexes. Heatmap with number of genes in different regulatory divergence categories for each expression inheritance profile to identify gene regulatory divergence underlying transgressive expression in autosomes. Males and females show different gene regulatory divergence underlying gene misexpression with higher contributions of *cis*-*trans* compensatory divergence to transgressive expression in males. The regulatory divergence for X-linked genes in males was inferred in a different way (see methods) due to the hemizygous nature of the X-chromosome in males (Supplementary Figure S2) but display similar pattern as autosomes and are depicted in Supplementary Figure S12. The top row shows results from H1 hybrids (*C. remanei* mother) and bottom row shows results from H2 hybrids (*C. latens* mother). **Alt text:** Grid of heatmaps mapping expression inheritance profiles against regulatory divergence categories. Top and bottom rows contrast H1 and H2 crosses across female soma, male soma, female gonad, and male gonad tissues, with shading density indicating gene counts and a blank space for H1 male gonad.

Transgressive expression provides a clear case of gene misexpression in hybrids that could underlie organismal dysfunction. Comparison of transgressive hybrid expression between the reciprocal crosses showed a higher contribution of compensatory *cis*-*trans* divergence for H1 than H2 in both sexes (p_MS_ = 0.01; p_FG_ = 0.0001; p_FS_ <0.001; Figure 6). Hybrid misexpression also could be argued to arise from an additive expression profile in between the two parents. While we detected additive expression effects relatively infrequently, it associates primarily with *cis-*only regulatory divergence (Figure 6), supporting the idea that *cis*-only divergence alone is a key mechanism of regulatory divergence between species.

### Expression inheritance and regulatory divergence in whole animals

Nearly all H1 males (*C. remanei* as the maternal parent) suffer from severe gonad developmental defects (Dey et al., 2014), and we were unable to reliably isolate gonad tissue for this class of males. To compensate for this challenge, we collected transcriptome data from male whole animal samples of *C. remanei*, *C. latens*, and hybrids from both cross directions (H1 and H2) to compare with soma tissue samples, in particular, which we hypothesized might reveal how inferences from whole male samples might be influenced by tissue allometric differences between H1 and H2.

In contrast to tissue-specific data, where H1 showed a slightly higher incidence of transgressive genes (Figure 2B, D), the whole male expression inferences imply more transgressive (underdominant and overdominant) expression for H2 than H1 (p < 0.001; n_transgressive, H1_ = 5584; n_transgressive, H2_ = 6142; n_total, H1_ = 13479; n_total, H2_ = 13479; Table 1; Supplementary Figure S13A, B). Whole male samples also implicated cross-dependent differences from male soma tissue in terms of regulatory divergence (Supplementary Figure S13C, D; Table 2; Table 3; cf. Figure 4B, D). Interestingly, however, approximately 68% of autosomal genes in whole male samples were inferred to have the same divergence category when assessed for male soma in the H1 cross (n=4255 genes out of 6264 total autosomal genes) (Supplementary Figure S14A). Similarly, around 70% of autosomal genes in H2 whole male samples showed the same divergence category when assessed for H2 male soma (n=4429 genes out of a total 6326 autosomal genes) (Supplementary Figure S14B). Whole male and soma tissue samples both showed significant enrichment of overdominant and additive genes on autosomes compared to the X-chromosome (Supplementary Figure S6); whole male samples were much more strongly enriched on the X-chromosome for *cis-trans* compensatory changes than were male soma tissue samples (Figure 5B; Supplementary Figure S14C, D). Comparison between whole male samples and male soma for X-linked genes, however, provides evidence for substantial sample-specific misregulation, as the overlap in genes in both hybrid crosses is just ∼30% despite identical X-chromosome origin (n_H1_ = 822; n_H2_ = 830 out of total 2614 X-linked genes; Supplementary Figure S15). The severe gonad developmental dysfunction in H1 males reduces its size and likely its cell count, creating tissue allometric differences between H1 and H2 in expression for whole male RNA samples due to different relative tissue compositions (Dey et al., 2014; Michalak & Noor, 2004). Consequently, as we observe here, misexpression in the gonad is likely to be more muted in whole male samples for H1 than for H2, leading to the counterintuitive result of more severe misexpression detected in whole male samples of H2 that successfully develop larger gonads with less severe tissue-scale dysfunction.

## Discussion

### Gene regulatory divergence and hybrid misexpression in nematodes

Gene regulatory divergence plays an important role in generating and maintaining species diversity by enabling adaptive phenotypic divergence and by genetically enforcing species boundaries. Specifically, divergent gene regulatory factors can cause hybrid dysfunction in interspecies hybrids due to the mismatch between diverged *cis*-acting regulatory elements and *trans*-acting factors from parental species (Landry et al., 2007; Mack & Nachman, 2017). We find extensive misexpression caused by gene regulatory divergence in the F1 hybrids of *C. remanei* and *C. latens*, with hybrid males displaying overall more genes subject to misregulation than hybrid females. More specifically, *cis*-*trans* compensatory divergence was pervasive in hybrid males, affecting approximately 8% of genes depending on tissue type and cross direction.

The high incidence of compensatory regulatory divergence suggests that *cis*-elements and *trans*-acting factors have frequently co-evolved to maintain conserved overall expression between *C. remanei* and *C. latens*. Stabilizing selection on expression phenotype is expected to often yield, in molecular evolutionary terms, such compensating regulatory changes that result in developmental system drift, in which divergence in developmental pathways occurs without observable changes in phenotype or expression (Cutter & Bundus, 2020; Gilad et al., 2006; Hodgins-Davis et al., 2015; True & Haag, 2001). Interestingly, we document sex biases in the incidence of compensatory *cis-trans* regulatory evolution. The prevalence of compensatory regulatory divergence in genes involved in male-biased genetic networks thus suggests a molecular evolutionary route for disproportionate male hybrid dysfunction, as evidenced in prior studies (Goncalves et al., 2012; Landry et al., 2005; Meiklejohn et al., 2003; Sánchez-Ramírez et al., 2021; Viswanath & Cutter, 2023). It remains unresolved, however, why stabilizing selection on male-biased genes might be predisposed to compensatory regulatory evolution whereas stabilizing selection on female-biased genes might be predisposed to regulatory conservation.

Mutations in *cis*-regulatory elements and *trans*-acting factors can result in differences in global gene expression between species, thereby giving rise to species-specific gene expression patterns (Mack & Nachman, 2017). Consistent with prior research, we infer a greater prevalence of *cis*-only relative to *trans*-only regulatory divergence based on patterns of allele-specific expression in the hybrids of *C. remanei* and *C. latens* (Figure 4B, D). The prevalence of genes with *cis*-only regulatory divergence may be attributed to the advantages of *cis*-regulatory mutations, which are often expected to exert fewer and less severe negatively-pleiotropic effects compared to *trans*-only mutations (Carroll, 2005; Stern & Orgogozo, 2009; Wray, 2007; Wray et al., 2003). While larger *trans*-acting allelic effects may contribute disproportionately to standing variation within species, such variants may not persist over the evolutionary long term or may foster subsequent *cis*-acting changes that compensate for negatively-pleiotropic effects (Coolon et al., 2015; Gomes & Civetta, 2015; Hill et al., 2021). The patterns we observe in *C. remanei* and *C. latens* are consistent with expectations that *cis*-regulatory elements play a disproportionate role in defining regulatory differences between species (Bell et al., 2013; Mack et al., 2016; Wittkopp et al., 2008).

Previous work with whole animal transcriptomic data for *C. briggsae* and *C. nigoni* nematodes characterized an even greater incidence of *cis*-*trans* compensatory changes than we documented for *C. remanei* and *C. latens* (Sánchez-Ramírez et al., 2021; Viswanath & Cutter, 2023). These species pairs, however, are distinct in multiple pertinent ways. *C. briggsae* and *C. nigoni* have a more ancient divergence and their hybrids show substantially more severe dysfunction than for *C. remanei* and *C. latens* (Dey et al., 2014; Woodruff et al., 2010; Fierst et al., 2015; Fusca et al., 2025). Moreover, *C. briggsae* and *C. nigoni* have divergent reproductive modes (androdioecy with self-fertilizing hermaphroditism vs gonochorism with obligatory mating) (Adams et al., 2023; Fierst et al., 2015; Yin et al., 2018), in contrast to the shared gonochorism of *C. remanei* and *C. latens*. The derived selfing hermaphroditism of *C. briggsae* has led to major genome rearrangements and gene losses, specifically with disproportionate loss of genes related to male function (Adams et al., 2023; Fierst et al., 2015; Yin et al., 2018). Consequently, we should anticipate the *C. briggsae*–*C. nigoni* system to differ both quantitatively and qualitatively from the findings for *C. remanei* and *C. latens*. Future investigations will prove insightful to determine the roles and relative contributions of divergence to additional mechanisms of gene regulation, including post-transcriptional regulators (e.g. via smallRNA pathways), alternative splicing, and protein regulation and how these factors might influence reproductive isolation.

### Sex-specific gene regulation and hybrid dysfunction

Sex-biased genes can experience distinct selective pressures in males and females within natural populations and often contribute to species differences due to faster evolution of male-biased genes, both in terms of coding sequence and expression levels, likely due to stronger sexual selection acting on males (Meiklejohn et al., 2003; Ranz et al., 2003; Parisi et al., 2004; Ellegren & Parsch, 2007; Assis et al., 2012; Harrison et al., 2015; Viswanath et al., 2025). Female-biased genes, on the other hand, often experience stronger evolutionary constraints as they contribute to roles with higher functional pleiotropy compared to male-biased genes, primarily because of their broader ranges of expression (Assis et al., 2012; Parisi et al., 2004; Presgraves & Meiklejohn, 2021).

Based on the potential for faster evolution of sex-biased genes and their contributions to species differences in expression (Viswanath et al., 2025), we hypothesized a disproportionate role for sex-biased gene misexpression in those male hybrids that experience more pronounced sterility. We found pervasive misexpression of sex-biased genes in hybrid males and females of *C. remanei* and *C. latens* (Figure 2E-H). Specifically, genes with transgressive expression in hybrid males tended to correspond to genes with male-biased or sex-neutral expression in parental species, with male-biased genes disproportionately showing underdominant expression in hybrid males. However, the misexpression in F1 females tended to masculinize their transcriptome relative to parental females by upregulating parentally-male-biased genes and downregulating parentally-female-biased genes (Figure 2E-H). Our analysis also revealed that gonads and soma have different sex-specific patterns of misexpression in the H2 cross, such that males show higher misexpression in gonads than females and that females have higher misexpression in the soma than males. Such transcriptional differences might manifest as a result of sexual selection acting differentially among tissues in males and females (Blekhman et al., 2008; Parisi et al., 2004; Small et al., 2009) and can have negative implications on sex-specific and/or reproductive processes such as disruption of spermatogenesis in males (Banho et al., 2021; Moehring et al., 2006; Ranz et al., 2004).

The F1 male hybrids of *C. remanei* and *C. latens* display pronounced hybrid sterility and hybrid inviability compared to females, demonstrating Haldane’s rule (Haldane, 1922; Turelli, 1998; Presgraves, 2010; Delph & Demuth, 2016; Mack & Nachman, 2017; Cowell, 2022; Cutter, 2024). The persistent misexpression of male-biased genes in hybrids of both sexes supports “faster male theory” for Haldane’s rule which proposes that dysfunction may arise from the rapid divergence of male-biased genes (Artieri & Fraser, 2014; Cowell, 2022; Cutter et al., 2019; Cutter & Ward, 2005; Sánchez-Ramírez et al., 2021). Interestingly, we find that the incidence of genes with transgressive misexpression in hybrid males is comparable to or lower than in females, for both gonad and soma tissues, despite more hybrid male dysfunction at the organismal level (Supplementary Figure S4 F, G). These observations contrast with some previous studies in *Drosophila* and mice (Haerty & Singh, 2006; Mack et al., 2016) but are similar to patterns seen in *Xenopus* frogs (Malone & Michalak 2008; Malone & Michalak 2008). Moreover, lower misexpression in hybrid males than females also was found for other *Caenorhabditis* species (*C. briggsae* × *C. nigoni*), suggesting a consistent difference in nematode biology compared to some other organisms (Sánchez-Ramírez et al., 2021). Possible explanations include greater absolute molecular divergence for nematode species pairs compared to flies or mice, as well as differences in nematode tissue and organismal development (Cutter, 2008; Fitch, 2005; Kiontke & Fitch, 2005; Mushegian et al., 1998; Riddle et al., 1997). We hypothesize that the more pronounced transgressive expression in hybrid females, despite elevated hybrid dysfunction in males, might indicate that the genetic networks of females are more resilient to perturbations in gene expression – and conversely that male genetic networks are more “fragile” – and thereby mitigate against organismal dysfunction in hybrid females (Meiklejohn et al., 2003; Michalak & Noor, 2004; Malone & Michalak, 2008b, 2008a; Gomes & Civetta, 2014, 2015; Harrison et al., 2015; R. Li et al., 2016; Sánchez-Ramírez et al., 2021).

Finally, we identified extensive *cis*-*trans* compensatory regulatory divergence from hybrid males for both reciprocal crosses (Figure 4B, D). These *cis*-*trans* compensating changes also contribute disproportionately to hybrid misexpression in males (Figure 4F, H) and show ∼7-fold enrichment on the X-chromosomes in males, whereas X-linked expression in females instead indicated enrichment of genes with conserved regulation. These patterns are consistent with stabilizing selection to preserve expression level acting on X-linked genes when expressed in both males and females. Interestingly, however, the molecular evolutionary outcomes of phenotypic stabilizing selection appear to depend on the sex in which expression takes place (Figure 5), with *cis*-*trans* compensatory regulatory divergence accruing disproportionately for male expression. Males and females can experience distinct selective pressures through sexual selection and to resolve sexual conflict, in addition to sex-specific natural selection. Such differential selection may contribute to the evolution of sex-biases in gene regulatory mechanisms in the form of *cis*-versus *trans*-acting regulatory controls. Furthermore, the abundance of male-specific *cis-trans* compensatory divergence, and greater rarity of genes with conserved regulation in males, provides a potentially important molecular evolutionary mechanism responsible for male-specific hybrid incompatibility at the organismal level, given that hybrid males are more severely affected by hybrid sterility and inviability than hybrid females. These observations suggest that sexual selection and stabilizing selection on gene expression might interact to contribute to sex-specific species differences in gene regulatory architectures that, in turn, contribute to disproportionate hybrid male dysfunction in the evolution of reproductive isolation between *C. remanei* and *C. latens*.

### Implications for Darwin’s corollary to Haldane’s rule

An interesting characteristic of *C. remanei × C. latens* hybrids is the extreme cross-dependent asymmetry in hybrid male sterility (Darwin’s corollary to Haldane’s rule; Turelli & Moyle, 2007), such that nearly all hybrid males with *C. remanei* mothers (H1) are sterile in large part due to gonad morphogenesis defects (Dey et al., 2012, 2014; Bundus et al., 2018; Dall’Acqua et al., 2025). Our analysis of hybrid males showed that asymmetries also manifest in terms of perturbed gene expression. Overall, gene misexpression is most evident in tissues of H1 animals, the hybrid cross displaying a greater degree of hybrid dysfunction at the organismal level. The 1.5-fold greater prevalence of genes with underdominant expression in H1 than H2 underlaid its elevated transgressive misexpression (Supplementary Figure S5B, C). Additionally, more than half of genes with underdominant expression in hybrid male soma samples were detected exclusively in just one of the hybrid cross directions, representing substantial cross-specific hybrid misexpression (Figure 3B, red circles). The X-chromosome, however, does not appear to play an outsized role in transgressive expression of hybrids (Supplementary figure S6).

The cross-dependent asymmetries in gene expression also reveal asymmetries in the inference of *cis*- and *trans*-acting regulatory divergence. Similar to patterns of gene misexpression, H1 hybrids revealed a consistently lower proportion of genes with conserved regulatory controls than H2 hybrids (Supplementary Figure S10A-C), with H1 male soma showing a nearly two-fold elevation in genes exhibiting *cis*-*trans* compensatory regulatory divergence. The *cis*-*trans* compensatory regulatory changes that act to conserve overall expression level of a given gene in males, however, are enriched on the X-chromosome.

Taken together, our observations demonstrate how diverged *cis*- and *trans*-acting regulatory factors can drive misexpression of genes in hybrid genomes in a cross-dependent manner. These profiles of asymmetric misexpression that are exacerbated in adult hybrid males that develop sterility is consistent with the “sterility hypothesis” that gene misexpression should accompany sterile hybrids (Gomes & Civetta, 2014, 2015). Moreover, X-linked genes experience disproportionate misregulation in hybrid males due to *cis*-*trans* compensatory divergence, consistent with a gene regulatory manifestation of the large X-effect (Coyne & Orr, 1998). Whole male transcriptomic data of hybrids between *C. briggsae* and *C. nigoni* also yield a similar large X-effect (Sánchez-Ramírez et al., 2021), and X-autosome hybrid incompatibilities have been documented in *Caenorhabditis* nematodes and could underlie the observed asymmetry in hybrid incompatibility (Bi et al., 2019; Bittorf, 2018; Bundus et al., 2018; Viswanath & Cutter, 2023). Although our approach constructively identified misregulated loci in hybrids due to *cis*-and *trans*-acting changes, which provide a group of candidate genes for testing a role in reproductive isolation, it remains a challenge from present data to determine conclusively whether the associated genes are causes of sterility or secondary consequences of defects in gonad morphogenesis or other aspects of prior development.

### Hybrid female paradox: regulatory asymmetry despite shared genomes

One feature of our analyses revealed an interesting conundrum regarding the patterns in hybrid females. Despite the fact that F1 hybrid females from both reciprocal cross directions have identical nuclear genome compositions, we detect *cis*-only divergence to be twice as prevalent in H1 females compared to H2 females (Supplementary Figure S10A, C). Furthermore, more than half of the autosomal genes demonstrating *cis*-*trans* compensatory regulatory divergence were distinct to H1, instead showing conserved regulation in H2 females, indicating that misregulation is exclusive to autosomal genes in H1 females that had *C. remanei* as the maternal parent (Figure 3D, F). Similar parent-of-origin effects have also been documented for autosomal genes in hybrid mice (Shen et al., 2014). Such asymmetries are counter-intuitive because hybrid females have identical nuclear genetic composition and, regardless of the cross direction, do not exhibit strong hybrid incompatibility at the organismal level (Dey et al., 2012, 2014).

Two ways that hybrid females differ in reciprocal crosses are in terms of their mitochondrial genomes and any intergenerationally inherited epigenetic factors (e.g., parentally-provisioned small RNAs or proteins), beyond the possibility of technical differences (e.g., statistical power) that might affect the ability to detect regulatory divergence. Thus, a possible resolution to the “hybrid female paradox” of cross-dependent differences is that genetic interactions involve mitochondria as cyto-nuclear incompatibilities (Ellison et al., 2008; Chang et al., 2016; Ågren et al., 2020; Nguyen et al., 2025; Sun & Yang, 2025). The existence of mito-nuclear incompatibilities is consistent with backcross analysis for *C. remanei* and *C. latens* (Bundus et al., 2018), and mito-nuclear epistatic interactions appear to be common within *C. elegans* (Nguyen et al., 2025).

Another possible resolution to the “hybrid female paradox” of cross-dependent differences is via the presence of epigenetic × nuclear interactions. Although *Caenorhabditis* lacks DNA methylation, several epigenetic pathways remain sensitive to disruptions in nematode hybrid genomes (Kelly, 2014; Rošić et al., 2018), including through maternally- or paternally-delivered transcription factor or other proteins, mRNA transcripts, small RNA regulators, or histone modifications (Brekke et al., 2021; Bundus et al., 2018; Burga et al., 2020; Kelly, 2014; Weiser & Kim, 2019). For example, incompatibility involving maternally-provided mismatches of histone variants such as H3.3 might contribute to dysfunction in hybrids, as such histone marks are crucial for chromatin reorganization during sperm decondensation in zygotes (Arico et al., 2011; Kelly, 2014; Rechtsteiner et al., 2010; Samson et al., 2014; Tabuchi et al., 2018). In addition to maternal provisioning, a number of paternally-delivered factors have been documented to influence embryos in *Caenorhabditis* (Seidel et al., 2008, 2011; Conine et al., 2013; Ben-David et al., 2017; Burga et al., 2020; Ben-David et al., 2021).

The expanded repertoire of small RNA pathways that nematodes have evolved can be delivered maternally or paternally to embryos (Conine et al., 2013, 2013; Weiser & Kim, 2019). The piRNA pathway in the germline of *C. elegans*, in particular, can mediate multigenerational silencing, act in a sex-specific manner, and can affect both spermatogenesis and oogenesis (Batista et al., 2008; Billi et al., 2013; Conine et al., 2010; Das et al., 2008; Seroussi et al., 2023; Tyc et al., 2017). Defects in piRNA pathways can also reactivate transposable elements, altering the epistatic landscape in hybrids, causing gene misexpression, reduced brood size, and fertility defects (Batista et al., 2008; Billi et al., 2013; Das et al., 2008; Serrato-Capuchina & Matute, 2018; Tyc et al., 2017). Furthermore, nematodes can employ RNAi-related mechanisms to identify foreign sequences based on a memory of past parental gene expression (Luteijn et al., 2012; Shirayama et al., 2012; Weiser & Kim, 2019). For example, maternally inherited RNAe can suppress paternally expressed transgenes in *C. elegans*, but endogenous genes are likely protected (Luteijn et al., 2012; Shirayama et al., 2012; Weiser & Kim, 2019).

Finally, another potential source of asymmetric misregulation involves selfish toxin–antidote (TA) elements. TA systems—identified in *C. elegans*, *C. tropicalis*, and *C. briggsae*—generate strong parent-of-origin effects through maternally or paternally deposited toxins (Ben-David et al. 2017, 2021; Seidel et al. 2011) that, in the absence of proper zygotic expression of an antidote, can impair embryonic or larval development, delay growth, or reduce hybrid fitness (Burga et al. 2020; Velazco-Cruz & Ross 2022). These TA systems often involve rapidly evolving, species-specific genes that also can intersect with small RNA pathway regulation (Pliota et al., 2024) and can contribute to asymmetric hybrid dysfunction in intra-specific hybrids and, potentially, interspecific hybrids (Ben-David et al., 2017, 2021; Burga et al., 2020; Seidel et al., 2008, 2011; Xie et al., 2026). While adverse TA effects would typically be expected in F2 and later generation hybrids, if the zygotic antidote does not get properly expressed in F1 embryos, then it becomes plausible that TA elements could contribute to hybrid incompatibility in the F1 generation.

Although mito-nuclear and epigenetic mechanisms can cause female asymmetry in hybrid misexpression, *C. remanei* × *C. latens* hybrid females show no substantial outward incompatibility phenotypes. Thus, the parent-of-origin effects on expression profiles that influence our inferences of regulatory divergence do not seem to cause major developmental problems for female hybrids. We hypothesize that the outputs of female genetic networks are more resilient to disruptions than are male gene regulatory networks, consistent with a “fragile male” view of hybrid dysfunction in Haldane’s rule (Wu & Davis, 1993; Cutter, 2024). While much current research emphasizes differences between sexes or between fertile versus sterile hybrids of a given sex, our findings underscore the additional value of characterizing classes of hybrids that do not exhibit significant hybrid incompatibility in order to gain a comprehensive understanding of the evolution and consequences of genic incompatibilities.

### Tissue composition in gene expression analysis

Differences in tissue composition can lead to differences in inferred gene expression profiles when using bulk RNA from heterogenous tissues such as whole bodies (Price et al., 2022). Such tissue allometric differences can be especially prominent in hybrids, given their propensity for dysfunctional development, and complicate the interpretation of differential gene expression (Harrison et al., 2015). This study reduces some of these concerns by dissecting distinct tissue samples, by focusing on closely related species that have very similar tissue size and composition between species, and by pooling tissues across many individuals for each replicate sample. The highly conserved deterministic cellular development in *Caenorhabditis*, and the exceptionally similar morphology and body size of *C. remanei* and *C. latens* that make them anatomically indistinguishable from one another (Dey et al., 2012, 2014), mitigates a direct influence of allometric effects. Nonetheless, the abnormalities in H1 male gonad development may still impact gene expression analysis and interpretation, especially when comparing gene expression from whole animals (Ranz et al., 2004; Sánchez-Ramírez et al., 2021; Sundararajan & Civetta, 2011). We indeed identified contrasting patterns of gene misexpression for whole male samples versus male soma alone using allele-specific expression analysis of hybrids, indicating that tissue allometric effects likely play a role in whole-animal analyses, supporting our choice to focus analysis on distinct dissected tissue types.

In whole male samples, we observed, counterintuitively, that males from the less-severely affected H2 cross exhibit a signal of more pronounced misexpression, whereas inferences from male soma tissue alone demonstrate the more intuitive higher prevalence of misexpression in H1 males that experience more severe developmental anomalies (Supplementary Figure S14A, B). The stronger developmental defects in H1 males likely affect the size, structure, cell count, and composition of the male gonad (Dey et al., 2014), leading to smaller gonads potentially causing allometric effects in whole male contrasts. If gonads are the primary source of misexpression in hybrid males, then the scarcity of misexpression in H1 whole male samples might simply reflect a reduced contribution of gonad transcripts to the whole male transcriptome due to their underdeveloped gonads. Previous research in *Drosophila* has also demonstrated contrasting gene expression patterns for whole animal and tissue-specific datasets, indicating false positives and a lack of ability to capture subtle tissue-specific effects (Catron & Noor, 2008; Haerty & Singh, 2006; Moehring et al., 2006; Sundararajan & Civetta, 2011). Despite whole animal experiments providing important first steps in deciphering gene regulatory evolution, especially in understudied hybrids and non-model organisms, these kinds of results emphasize the value of incorporating tissue-specific expression when possible (Catron & Noor, 2008; Montgomery & Mank, 2016; Price et al., 2022).

## Acknowledgements

This work was supported by funds from the Natural Sciences and Engineering Research Council of Canada to A.D.C. and J.A.C. We thank Yunchen Gong from the University of Toronto’s Centre for the Analysis of Genome Evolution and Function (CAGEF) for generating the updated annotations for *C. remanei* and *C. latens*. We are grateful to Trisha Wittkopp and the members of the Cutter lab for their helpful and constructive feedback on this work.

---

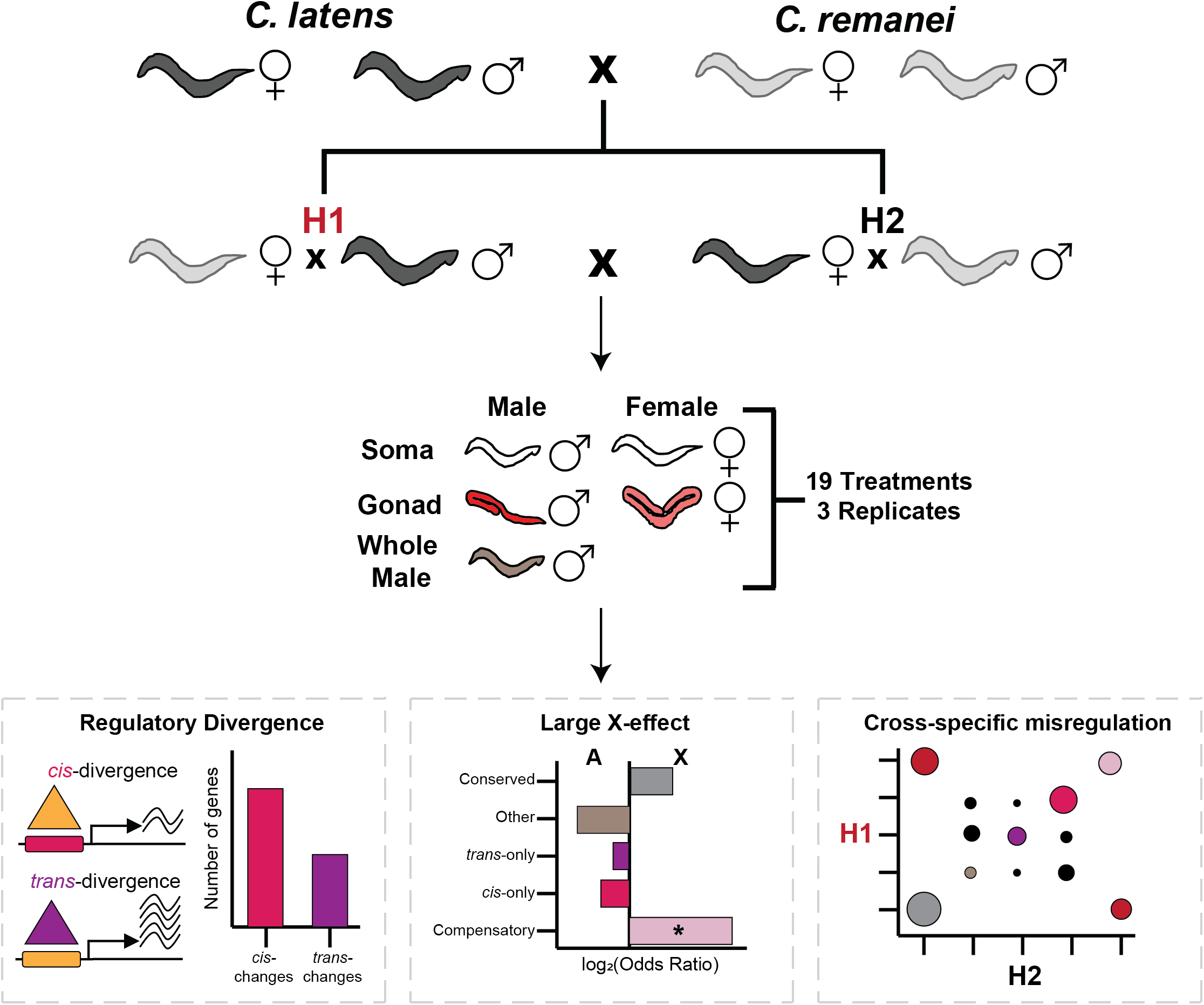

